# Benchmarking Orientation Distribution Function Estimation Methods for Tractometry in Single-Shell Diffusion Magnetic Resonance Imaging - An Evaluation of Test-Retest Reliability and Predictive Capability

**DOI:** 10.1101/2025.09.02.673635

**Authors:** Amelie Rauland, Steven L. Meisler, Aaron F. Alexander-Bloch, Joëlle Bagautdinova, Erica B. Baller, Raquel E. Gur, Ruben C. Gur, Audrey C. Luo, Tyler M. Moore, Oleksandr V. Popovych, Kathrin Reetz, David R. Roalf, Russell T. Shinohara, Susan Sotardi, Valerie J. Sydnor, Arastoo Vossough, Simon B. Eickhoff, Matthew Cieslak, Theodore D. Satterthwaite

## Abstract

Deriving white matter (WM) bundles in-vivo has thus far mainly been applied in research settings, leveraging high angular resolution, multi-shell diffusion MRI (dMRI) acquisitions that enable advanced reconstruction methods. However, these advanced acquisitions are both time-consuming and costly to acquire. The ability to reconstruct WM bundles in the massive amounts of existing single-shelled, lower angular resolution data from legacy research studies and healthcare systems would offer much broader clinical applications and population-level generalizability. While legacy scans may offer a valuable, large-scale complement to contemporary research datasets, the reliability of white matter bundles derived from these scans remains unclear. Here, we leverage a large research dataset where each 64-direction dMRI scan was acquired as two independent 32-direction runs per subject. To investigate how recently developed bundle segmentation methods generalize to this data, we evaluated the test-retest reliability of the two 32-direction scans, of WM bundle extraction across three orientation distribution function (ODF) reconstruction methods: generalized q-sampling imaging (GQI), constrained spherical deconvolution (CSD), and single-shell three-tissue CSD (SS3T). We found that the majority of WM bundles could be reliably extracted from dMRI scans that were acquired using the 32-direction, single-shell acquisition scheme. The mean dice coefficient of reconstructed WM bundles was consistently higher within-subject than between-subject for all WM bundles and ODF reconstruction methods, illustrating preservation of person-specific anatomy. Further, when using features of the bundles to predict complex reasoning assessed using a computerized cognitive battery, we observed stable prediction accuracies (*r*: 0.15-0.36) across the test-retest data. Among the three ODF reconstruction methods, SS3T had a good balance between sensitivity and specificity in external validation, a high intra-class correlation of extracted features, more plausible bundles, and strong predictive performance. More broadly, these results demonstrate that bundle segmentation can achieve robust performance even on lower angular resolution, single-shell dMRI, with particular advantages for ODF methods optimized for single-shell data. This highlights the considerable potential for dMRI collected in healthcare settings and legacy research datasets to accelerate and expand the scope of WM research.

## 1 Introduction

Diffusion-weighted magnetic resonance imaging (dMRI) measures the directional diffusion of water molecules and is the key modality for non-invasively estimating white matter (WM) microstructural tissue properties (Alexander et al., 2019; Le Bihan et al., 2001) and reconstructing WM fiber pathways using tractography algorithms (Jeurissen et al., 2019; F. Zhang et al., 2022). Advanced post-processing methods that can model multiple compartments (Assaf & Basser, 2005; H. Zhang et al., 2012) or multiple fiber populations (Jeurissen et al., 2014) per voxel improve reconstructions of fiber pathways and microstructural tissue properties. However, these methods require high-angular resolution data acquired on multiple diffusion shells as is common in large-scale neuroimaging datasets like the Human Connectome Project (Van Essen et al., 2013), UK Biobank (Bycroft et al., 2018), or the ABCD study (Garavan et al., 2018) with up to 270 directions acquired on up to 4 shells. Such scans are very time- and resource-intensive to acquire, limiting their practicality to populations less tolerant to MRI. In contrast, the overwhelming majority of dMRI data acquired over the past 30 years, as part of research protocols or in healthcare systems, is from scans with fewer directions (between 6 and 32) and only a single-shell (typically *b* < 1200 s/mm²). In this work, we refer to such single-shell protocols with lower angular resolution as *simple acquisitions* and the aforementioned multi-shell, higher angular resolution protocols as *advanced acquisitions*. We sought to investigate how reliably WM bundles can be reconstructed from these simple acquisitions and assess their utility for establishing brain-behavior relationships. Understanding the reliability of such legacy or clinically acquired data is crucial, as prediction analyses of inter-individual differences benefit from large sample sizes (Cui & Gong, 2018; Gell et al., 2024; Scheinost et al., 2019).

Important gaps remain in understanding the reliability of reconstructions derived from single-shell, low angular resolution scans. For instance, it has been shown that lower angular resolution affects WM bundle reconstruction (Ambrosen et al., 2020; Calabrese et al., 2014; Radhakrishnan et al., 2023; Vos et al., 2016; Wasserthal et al., 2018) and derived microstructural properties (Aja-Fernández et al., 2023; Lebel et al., 2012; Spagnolo et al., 2024; Zhan et al., 2010). These comparisons typically highlight differences between simple and advanced acquisitions, with greater discrepancies arising from larger differences in acquisition protocols. While these studies primarily demonstrate that simple acquisitions often yield less accurate results than advanced scans with multiple shells and a high angular resolution, they leave open the important question of whether simple scans can produce metrics that are sufficiently reliable for research use.

Following image acquisition, the choice of the orientation distribution function (ODF) reconstruction algorithm has been shown to impact the resulting WM microstructural properties and bundle segmentations (Daducci et al., 2014; Sydnor et al., 2018; Tournier et al., 2012; Xie et al., 2015). In a study using single-shell scans, it has been shown that different methods for reconstructing the ODF affect the characteristics of reconstructed WM bundles, such as their completeness and the presence of false positives (Wilkins et al., 2015). However, as the study included only one scan per subject, it was impossible to evaluate the impact of the ODF reconstruction method on the test-retest reliability of the reconstructed tracts.

Finally, it remains unclear whether single-shell scans with low angular resolution are suitable for establishing robust links between brain structure and behavior, as most studies investigating such associations rely on high-quality, densely sampled diffusion data (Dhamala et al., 2021; Lo et al., 2025; Ooi et al., 2022; Yeung et al., 2023). Evidence from microstructural analyses suggests that simpler acquisitions may compromise sensitivity to meaningful effects. For example, Aja-Fernández et al. (2023), found that decreasing diffusion directions from 61 to 21 altered diffusion tensor imaging (DTI) metrics and hindered the detection of group differences between episodic and chronic migraine patients in a sample of 100 migraine patients (Aja-Fernández et al., 2023). Assessing brain-behavior associations in large sample sizes using WM bundle features extracted from such simple scans is therefore critical to determine whether such data can support studying individual differences.

Here, we investigated whether single-shell diffusion MRI scans with limited angular resolution could reliably segment person-specific WM bundles and support analyses of brain-behavior relationships. We also examined whether different ODF reconstruction methods systematically affected the test-retest reliability, overlap with advanced atlas bundles, and predictive performance. To address these questions, we used data from the Philadelphia Neurodevelopmental Cohort (PNC, (Satterthwaite et al., 2014)), a large-scale research dataset where each 64-direction dMRI scan was acquired as two independent 32-direction runs per subject, which are similar to scans acquired by some healthcare systems and many legacy research datasets. As described below, our results showed promising test-retest reliability for the majority of reconstructed WM bundles, high correspondence with atlas bundles derived from advanced acquisitions, and stable accuracy when predicting cognition from features of the reconstructed bundles, warranting the use of legacy research data or healthcare system images for research.

## 2. Materials and Methods

### 2.1 Participants

This study utilized data from the Philadelphia Neurodevelopmental Cohort (PNC) (Satterthwaite et al., 2014), a community-based sample of children, adolescents, and young adults aged 8-21 from the greater Philadelphia area, designed to investigate brain development. The cohort includes psychiatric and cognitive phenotyping for 9,428 participants, with a subsample of 1,445 individuals who underwent multimodal neuroimaging. For our analysis, we used each participant’s structural T1-weighted (T1w) scan, DWI data, and scores from the Penn Computerized Neurocognitive Battery (Gur et al., 2012), which assesses a range of cognitive domains. All participants over the age of 18 provided written informed consent prior to participation. For individuals under 18, informed assent was obtained along with written parental consent, and all participants received monetary compensation. The study was approved by the Institutional Review Boards of the University of Pennsylvania and the Children’s Hospital of Philadelphia.

### 2.2 Neuroimaging Acquisition

All MRI scans were acquired on the same 3T Siemens Tim Trio scanner (software version VB17) with a 32-channel head coil at the Hospital of the University of Pennsylvania (Satterthwaite et al., 2014).

The T1-weighted structural images were acquired using a magnetization-prepared rapid acquisition gradient-echo (MPRAGE) sequence with the following set of parameters: Repetition time (TR) = 1,810 ms, echo time (TE) = 3.51 ms, inversion time = 1,100 ms, flip angle = 9°, field of view (FOV) = 180×240 mm, matrix = 192×256, number of slices = 160, and the voxel resolution = 0.94×0.94×1 mm.

Diffusion scans were acquired using a distortion-minimizing twice-refocused spin-echo (TRSE) single-shot echo-planar imaging (EPI) sequence with the following set of parameters: TR = 8100ms, TE = 82 ms, FOV = 240x240 mm, matrix = 128x128, number of slices = 70, slice thickness/gap = 2/0 mm, flip angle = 90/180/180, and voxel resolution = 1.875×1.875×2mm. The total DWI acquisition protocol included 64 diffusion-weighted volumes with a *b*-value of 1000 s/mm^2^ along with 7 non-diffusion-weighted (*b*=0 s/mm^2^) volumes. To reduce the scan time for study participants, the acquisition was split into two separate scans, each with 32 diffusion encoding directions independently sampling the sphere that were acquired within the same hour. To be combined into a single 64-direction scan, the gradient schemes differed between the two 32-direction scans (exact *b*-vectors for each scan can be found in (Satterthwaite et al., 2014)). The first scan had 3 *b*=0 images, and the second had 4 *b*=0 images. Here, we did not combine the two scans into a single 64-direction scan. Instead, we considered them as independent, clinically feasible acquisitions, enabling an assessment of test-retest reliability.

### 2.3 Sample Construction

The complete sample construction procedure is depicted in **Figure 1**. We started with a total of *n* = 1,397 subjects, which had both DWI scans and the structural T1w image available. One subject was excluded due to unsuccessful processing, and 12 subjects were excluded due to deviation from the acquisition protocol (e.g., varied repetition time or voxel size). Missing fieldmaps were not considered a disqualifying acquisition variant because fieldmaps were not leveraged during the preprocessing of the data to better align with a typical clinical DWI acquisition protocol. Of the resulting subjects, 163 were excluded based on the quality control (QC) performed in (Roalf et al., 2016). This led to a total of n = 1,221 subjects that were included in the reliability analysis of reconstructed WM bundles.

**Figure 1:**
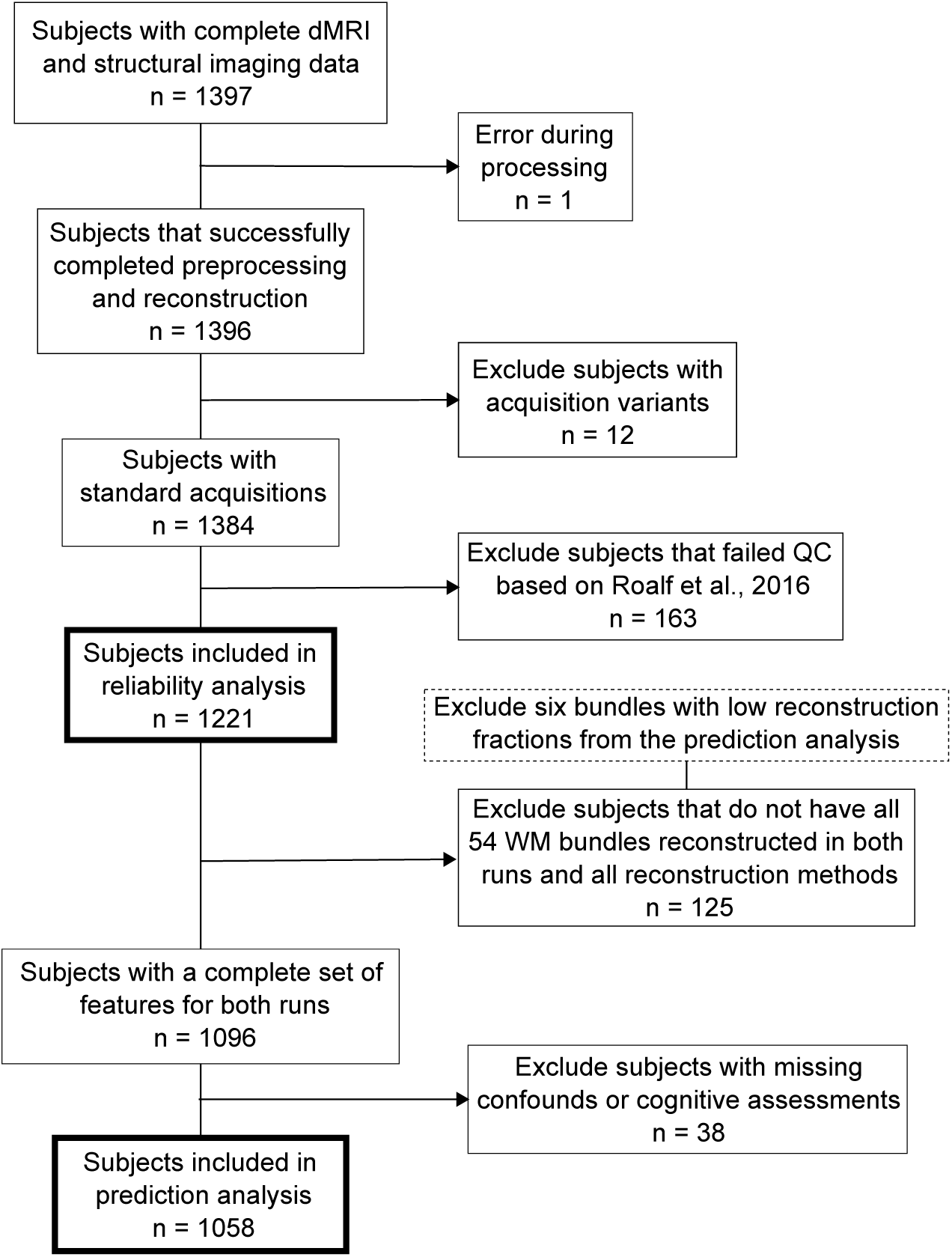
Subject inclusion flowchart illustrating the derivation of the final sample sizes for the reliability and prediction analysis.

For the prediction analysis, 6 out of 60 WM bundles were excluded because of low bundle reconstruction success rates (see below). The prediction analysis included subjects for which all 54 resulting WM bundles were reconstructed for both DWI scans for all three ODF reconstruction methods, leading to an exclusion of 125 subjects. Further, 38 subjects were excluded based on missing covariates, such as cognitive assessments required for prediction. The final number of subjects included in the main prediction analysis was n = 1,058.

### 2.4 Data Processing and Bundle Reconstruction

**Figure 2** illustrates the data processing workflow. The raw diffusion data were minimally pre- processed before undergoing three different methods for ODF reconstruction applicable to single-shell data with low angular resolution, and subsequent WM bundle tractography based on the ODFs.

**Figure 2:**
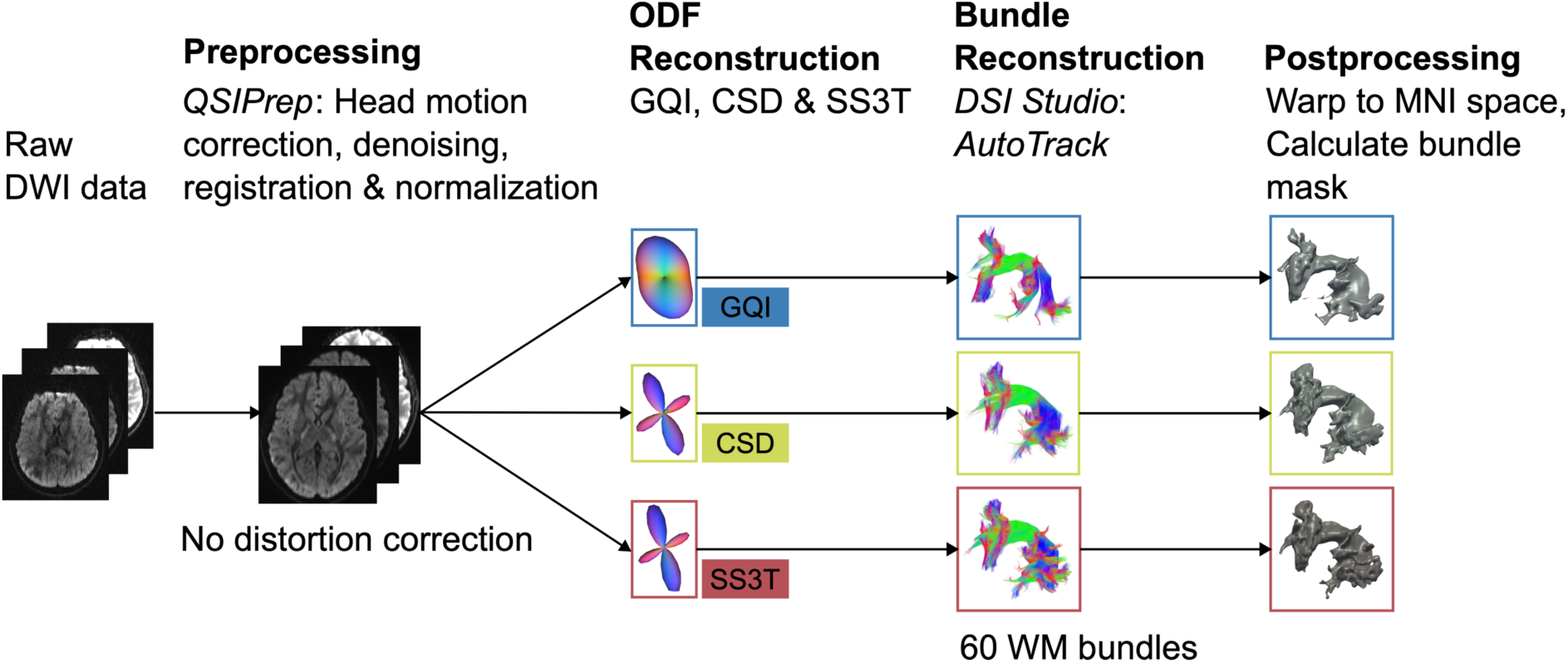
Schematic describing the DWI processing pipeline. After preprocessing the raw DWI data, orientation distribution functions (ODF) were reconstructed using three different methods suitable for single-shell data. Subsequently, 60 known WM bundles were extracted based on the ODFs using *DSI Studio AutoTrack*. The resulting bundles were then warped to MNI space and transformed from streamlines to a 3D binary voxel-wise mask for further analysis. Exemplary ODFs were derived from the same WM voxel across methods. The illustrated exemplary WM bundle is the left arcuate fasciculus. The streamline and bundle visualizations were created using *MI-Brain* (Rheault et al., 2016).

#### 2.4.1 Preprocessing

Both DWI scans and the T1w scan were pre-processed using *QSIPrep* 0.21.4 (Cieslak et al., 2021). Some of the text in the following section is boilerplate text that was automatically generated in *QSIPrep,* released under the CC0 license for reuse in manuscripts.

##### 2.4.1.1 Anatomical Preprocessing

The T1w image was corrected for intensity non-uniformity (INU) using N4BiasFieldCorrection (Tustison et al., 2010), *ANTs* 2.4.3), and used as an anatomical reference throughout the workflow. The anatomical reference image was reoriented into AC-PC alignment via a 6-DOF transform extracted from a full Affine registration to the MNI152NLin2009cAsym template. A full nonlinear registration to the template from the AC-PC space was estimated via symmetric nonlinear registration (SyN) using *antsRegistration*. Brain extraction was performed on the T1w image using *SynthStrip* (Hoopes et al., 2022), and automated segmentation was performed using *SynthSeg* (Billot, Greve, et al., 2023; Billot, Magdamo, et al., 2023) from *FreeSurfer* version 7.3.1.

##### 2.4.1.2 DWI Preprocessing

The two DWI scans per subject were preprocessed completely independently: Any images with a *b*-value less than 100 s/mm^2^ were treated as a *b*=0 image. MP-PCA denoising, as implemented in *MRtrix3*’s *dwidenoise* (Veraart et al., 2016), was applied with a 5-voxel window. After MP-PCA, the mean intensity of the DWI series was adjusted so that all the mean intensities of the *b*=0 images matched across each separate DWI scanning sequence. B1 field inhomogeneity was corrected using *dwibiascorrect* from *MRtrix3* with the N4 algorithm (Tustison et al., 2010) after corrected images were resampled.

*FSL’s* (version 6.0.7.9) *eddy* was used for head motion correction and *Eddy* current correction (Andersson & Sotiropoulos, 2016). *Eddy* was configured with a *q*-space smoothing factor of 10, a total of 5 iterations, and 1000 voxels used to estimate hyperparameters. A linear first-level model and a linear second-level model were used to characterize Eddy current-related spatial distortion. *q*-space coordinates were forcefully assigned to shells. Field offset was attempted to be separated from subject movement. Shells were aligned post-eddy. *Eddy*’s outlier replacement was run (Andersson et al., 2016). Data were grouped by slice, only including values from slices determined to contain at least 250 intracerebral voxels. Groups deviating by more than 4 standard deviations from the prediction had their data replaced with imputed values. Final interpolation was performed using the *jac* method.

The framewise displacement (FD) was calculated based on the preprocessed DWI using the implementation in *Nipype* (following the definitions by (Power et al., 2014)). The DWI time series were reoriented to AC-PC, generating a preprocessed DWI run in AC-PC space with 1.7mm isotropic voxels.

While fieldmaps are available in the PNC dataset, they were not used for distortion correction to better emulate clinical acquisition conditions. In routine clinical settings, diffusion-weighted imaging is often performed without acquiring fieldmaps, and distortion correction based on fieldmaps would therefore not be possible for these types of scans.

#### 2.4.2 ODF and Bundle Reconstruction

Following preprocessing of the structural and diffusion-weighted data, we reconstructed ODFs, which serve as the basis for tractography and subsequent WM bundle reconstruction. We compared three commonly used ODF reconstruction methods suitable for single-shell, low angular resolution data:

- Generalized *q*-Sampling Imaging (GQI) (F.-C. Yeh et al., 2010) is a model-free approach that estimates spin distribution functions (SDFs) directly from the diffusion MRI signals using a Fourier transform relation. This enables direct voxel-wise comparisons and can be applied to both grid and shell sampling schemes.
- Constrained spherical deconvolution (CSD) (Tournier et al., 2007) estimates a fiber ODF (fODF) by deconvolving the measured signal with a response function - an estimate of the signal expected from a single-fiber WM population. However, because CSD models only the WM signal, it may produce distorted FODs in voxels affected by partial voluming with gray matter (GM) or cerebrospinal fluid (CSF).
- Single-shell three-tissue CSD (SS3T) (Dhollander & Connelly, 2016) builds on the CSD algorithm by estimating separate WM, GM, and CSF signal components. While multi-tissue CSD traditionally requires multi-shell data (Jeurissen et al., 2014), SS3T achieves similar separation using only single-shell (+*b*=0) data via a specialized optimization algorithm.

We also used *DSI Studio* to generate maps of quantitative anisotropy (QA, (F.-C. Yeh et al., 2010)), isotropic diffusion (ISO), and DTI-derived scalars. Using the reconstructed ODFs, we attempted to reconstruct 60 known WM bundles (**Supplementary Table S1**) with the *AutoTrack* algorithm implemented in *DSI Studio*. *AutoTrack* is based on a tractography atlas (F.-C. Yeh et al., 2018) that uses a QA+ISO-based non-linear registration to each subject’s native diffusion space. For each WM bundle, seed points for deterministic tractography are placed within the corresponding atlas-defined bundle region in the subject space. Generated streamlines are retained only if they are sufficiently similar to an atlas streamline based on the Hausdorff distance. For each successfully reconstructed bundle, shape metrics (F.-C. Yeh, 2020) as well as diffusion tensor imaging (DTI) metrics were extracted from the resulting segmentation.

All three ODF reconstruction methods and bundle reconstruction were performed using *QSIRecon* 0.23.2. For GQI, we used the *DSI Studio* Hou version, with a mean diffusion distance ratio of 1.25. CSD reconstruction was run using *MRtrix3* (Tournier et al., 2019) based on single-fiber response functions estimated using the *tournier* algorithm (Tournier et al., 2013). For SS3T, multi-tissue fiber response functions were estimated using the *MRtrix3 dhollander* algorithm (Dhollander et al., 2019), and SS3T-CSD was performed using *MRtrix3Tissue* (https://3Tissue.github.io), a fork of *MRtrix3* (Tournier et al., 2019). Both CSD and SS3T FODs were intensity-normalized using *mtnormalize* (Raffelt et al., 2017). *AutoTrack* parameters were as follows: The distance tolerance was set to 22, 26, or 30 mm (evaluating all three and selecting one per bundle), the track-to-voxel ratio to 2.0, and the yield rate to 1.0e-06. While ODF maps generated with GQI are the standard input to the *AutoTrack* algorithm, the ODF maps generated using CSD and SS3T had to be converted using the *mif2fib* conversion implemented in *QSIRecon* to be used as input to *AutoTrack*.

Finally, all the reconstructed bundles were warped from the subject space to the MNI152NLin2009cAsym space. This was implemented by leveraging the transformation generated by *QSIPrep* to warp between the subject AC-PC and MNI space during preprocessing and adapting it such that it can be applied to the streamlines in *MRtrix3*. A binary mask image was created where voxels traversed by at least one streamline were set to one. Standardizing bundles in MNI space allowed us to assess the spatial alignment across individuals and coverage provided by the bundles.

### 2.4 Reliability Analysis

The reliability analysis was conducted in several steps, each described in detail below.

#### 2.5.1 Bundle Reconstruction Success Rate

First, we evaluated the reconstruction success rate for each bundle across all participants and DWI sessions. It was calculated as the number of successful reconstructions divided by the total number of scans collected over all subjects. We consider a reconstruction successful if an output file was generated with reconstructed streamlines for the bundle of interest. If no streamlines were found that were sufficiently similar to the atlas bundle of interest in terms of Hausdorff distance, the *AutoTrack* algorithm did not return an output for that respective bundle and we consider the reconstruction of that bundle for this scan to be unsuccessful. This step does not yet consider the quality of the reconstruction; it only describes whether a given bundle could be reconstructed at all.

#### 2.5.2 Dice Scores

In the second step of the reliability analysis, the Dice overlap between any two reconstructions of the same bundle was calculated. The Dice overlap is a common metric in segmentation tasks, probing the overlap between the ground truth (GT) and generated segmentation. It is defined as in **Eq. 1**, where *X* and *Y* are two bundle masks from different scans.

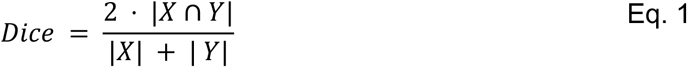

It can take values between 0, no overlap, and 1, perfect overlap. Here, we evaluated whether the Dice score of a given bundle is higher for two scans of the same subject (i.e., within-subject similarity) compared to any two scans of different subjects (i.e., across-subject similarity). For each bundle and ODF reconstruction method, we computed 1221 within-subject Dice scores (one per subject) and 2,979,240 between-subject Dice scores. We arrived at the number of between-subject Dice scores as described in **Eq. 2**, where *n* is the number of subjects.

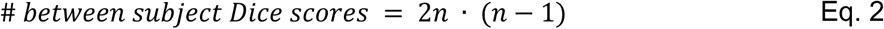

This corresponds to all four possible pairings of scans across two different individuals - that is, both runs from each subject were compared with both runs from the other subject. If a bundle was not reconstructed in one or both of the scans being compared, the corresponding Dice score was set to NaN. This differentiates missing reconstructions from cases where the bundle was reconstructed in both scans but showed no spatial overlap (Dice = 0).

#### 2.5.3 Discriminability

Discriminability is a statistical measure of test-retest reliability that quantifies how well repeated measurements from the same subject can be distinguished from measurements across different subjects (Bridgeford et al., 2021; Wang et al., 2024). It is defined as the proportion of times that the within-subject distance is smaller than the between-subject distance. A discriminability score of 1 is the highest possible score and indicates that for every subject, the distance between their two images is smaller than the distance between that subject’s images and any image from a different subject.

In our analysis, we defined the distance between two bundle reconstructions as 1 − Dice. Using these distances, we computed discriminability separately for each of the 60 white matter bundles using the *hyppo* Python package (Panda et al., 2024). Further, we compared the discriminability values for the same bundle between different ODF reconstruction methods. To assess whether discriminability differed significantly between the three ODF reconstruction methods per bundle, we used permutation tests implemented in the *hyppo* Python package (Panda et al., 2024). To assess whether discriminability differed significantly between the overall distributions including all bundles, we used the Wilcoxon signed-rank test (Wilcoxon, 1992) and corrected for multiple comparisons using the Benjamini-Hochberg procedure (Benjamini & Hochberg, 1995). For each bundle, only subjects for whom the bundle was successfully reconstructed in both scans across all three methods were included in the analysis.

#### 2.5.4 Comparison to Atlas Bundles

While discriminability indicates whether a bundle reconstruction is more similar within a subject than between subjects, it does not assess the biological plausibility of the reconstruction. To this end, we implemented an external validation by comparing the reconstructed bundles to the corresponding atlas bundles, which were derived from advanced diffusion data and curated by a team of neuroanatomists (F.-C. Yeh et al., 2018). To do this, the atlas bundles were masked and warped from the MNI152NLin2009bAsym to the MNI152NLin2009cAsym space using a non-linear transform with *antsApplyTransforms* (Avants et al., 2009) to be in the same space as the reconstructed bundles. For each bundle reconstruction, we calculated sensitivity and specificity by comparing the reconstructed bundle to the corresponding atlas bundle. Here, a voxel was considered a true positive if it was present in both the atlas and our reconstruction, a false negative if it was in the atlas but not in the reconstruction, a true negative if it was in neither, and a false positive if it was not part of the atlas bundle but present in the reconstruction. To avoid inflated specificity values due to the large background volume compared to the bundle volume, we limited the calculation to voxels that belonged to the union of the atlas bundle and all reconstructed versions of the bundle.

Additionally, probabilistic population maps were created for each bundle by adding up all masks of the reconstructed bundle and dividing by the total number of reconstructions. These maps, with values between [0, 1], reflect the frequency with which each voxel was assigned to the bundle. A voxel with a value of zero indicates it was never part of the bundle, while a value of one means it was included in every reconstruction. Visualizing these maps overlaid on the atlas bundles provided an intuitive assessment of anatomic coverage and reconstruction completeness for each ODF method.

#### 2.5.5 Feature Reliability

While the reliability of bundle shape, location, and extent is crucial for subsequent analyses, the bundles themselves are typically not used directly in brain-behavior studies. Instead, scalar features extracted from these bundles, such as mean diffusivity (MD), fractional anisotropy (FA), and bundle volume, serve as inputs for predictive modeling. To link bundle reliability and prediction performance, it is therefore important to also assess feature reliability. To this end, we calculated the intraclass correlation coefficient (ICC) for each bundle and ODF reconstruction method for the three features considered in our prediction analysis. For every feature, such as a particular bundle’s volume, we obtained two values per ODF reconstruction method, one from each of the two diffusion scans. Although these values were treated separately during prediction (see Section 2.6), we used them jointly to estimate test-retest reliability. Specifically, we computed ICC values for bundle volume, mean FA, and mean MD for all 60 bundles and across all three ODF reconstruction methods. The ICC(1,1), as defined by Shrout and Fleiss (Shrout & Fleiss, 1979), was calculated using the *pingouin* package (Vallat, 2018). ICC values generally range from 0 to 1 and can be categorized into four levels of test-retest reliability: excellent (ICC > 0.75), good (ICC = 0.60 to 0.74), fair (ICC = 0.40 to 0.59), and poor (ICC < 0.40) (Fleiss et al., 2003). It should be noted that the ICC was calculated for all 60 WM bundles for completeness. The final prediction analysis, however, only included 54 out of the 60 bundles (see Section 2.6.1).

To test for significant differences between ICC distributions of different ODF reconstruction methods, we used the Wilcoxon signed-rank test (Wilcoxon, 1992) and corrected for multiple comparisons using the Benjamini-Hochberg procedure (Benjamini & Hochberg, 1995).

### 2.6 Prediction Analysis

After assessing the reliability and completeness of the reconstructed bundles, we conducted a second main analysis to evaluate whether features derived from these bundles can predict inter-individual differences in cognition.

#### 2.6.1 Prediction Framework

To assess the ability of features extracted from the reconstructed bundles to predict inter-individual differences, we extracted features from 54 WM bundles for use in the prediction task. Six bundles - optic radiation (L/R), corticobulbar tract (L/R), and dentatorubrothalamic tract (left-to-right/right-to-left) - were excluded due to low reconstruction success rates and consequently frequent NaN values in the extracted features. The features considered for each bundle were the bundle volume (mm^3^), mean FA, and mean MD. This resulted in four different feature sets: (1) all three features for all bundles (162 features total), and (2)-(4) each individual feature (volume, FA, or MD) across all bundles (54 features each).

As the prediction target, we used a complex reasoning accuracy score from the Penn Computerized Neurocognitive Battery (PCNB, (Gur et al., 2010, 2012; Moore et al., 2015)). The PCNB includes 14 cognitive tests adapted from functional neuroimaging paradigms to assess various cognitive domains. In the main analysis, we focused on a summary score for complex reasoning (verbal reasoning, nonverbal reasoning, and spatial processing). In a sensitivity analysis, we considered two additional targets from the PCNB, to ensure findings can be replicated across targets (see Section 2.6.4).

We employed a linear ridge regression model for prediction. A nested 5-fold continuous stratified cross-validation (CV) was used, with 100 repetitions to estimate the distribution of prediction performance. Using a stratified splitting approach ensures that the distributions of the target variable are similar in the training and test sets. The regularization parameter α was tuned within the inner folds of the nested CV.

To account for potential confounding variables, we regressed out participant age (in months), mean framewise displacement (FD) (as output by *QSIPrep*), and sex from the features in a CV-consistent manner. Features were standardized (z-scored) before model training.

For each of the four feature sets, we conducted predictions separately for each of the three ODF reconstruction methods (GQI, CSD, SS3T) and both runs. This yielded a total of 24 distinct prediction pipelines in the main analysis (4 feature sets x 3 methods x 2 runs). The prediction pipeline, including confound removal, feature preprocessing, and CV splits, was implemented using *Julearn* (Hamdan et al., 2024), which is based on *scikit-learn* (Pedregosa et al., 2011).

#### 2.6.2 Prediction Accuracy

Prediction accuracy was assessed by calculating the Pearson correlation coefficient (*r*) between the observed and predicted cognition scores on the test set. We derived a distribution of 500 correlation values for each framework, corresponding to 100 repetitions of the 5-fold CV, resulting in 500 test sets from which correlations were calculated. In the main analysis, which focused on predicting complex reasoning, we tested for significant differences in prediction accuracy between the three ODF reconstruction methods under otherwise identical conditions, i.e., same scan and same feature group (e.g., all features (volume, MD and FA) from the CSD reconstruction of the first scan vs. all features from the SS3T reconstruction of the first scan). To assess these differences, we applied a specialized *t*-test designed for comparing machine learning algorithms, which accounts for variability not only from the choice of the test set but also from the training set (Nadeau & Bengio, 2003). The implementation of this *t*-test was provided in *Julearn* (Hamdan et al., 2024). The resulting *p*-values were further corrected for multiple comparisons using the Benjamini-Hochberg procedure (Benjamini & Hochberg, 1995). For the main analysis, we further evaluated the mean squared error (MSE), as a proxy for prediction accuracy.

#### 2.6.3 Prediction Similarity

For each combination of feature group and ODF reconstruction method, we compared predictions made from features of the first run and the second run. Ideally, predictions should be very similar across runs. To assess this similarity, we calculated the Pearson correlation between the predicted scores from the first run and those from the second run across all 500 test sets. This procedure provided a distribution of Pearson correlations, which we then compared between different ODF reconstruction methods and feature groups. To ensure a valid comparison, we used the exact same cross-validation splits for both runs, so that predictions from the first and second scan were generated for the same individuals in each fold. This was necessary to avoid correlating predictions from different subject groups, which would not reflect the stability of prediction across scans for the same individuals.

#### 2.6.4 Sensitivity Analysis

To investigate whether our results can be replicated for other prediction frameworks, we further investigated the prediction of two additional targets and one additional confound. For the two targets, we considered the summary score for executive functioning (abstraction and mental flexibility, attention, and working memory) and an overall accuracy score over all tests included in the PNCB. As an additional confound, we added total brain volume (TBV) to the original set of confounds (sex, age, mean FD).

## 3. Results

We evaluated the potential of WM bundles reconstructed from single-shell DWI data with a limited angular resolution of 32 directions through two main analyses. First, we assessed the reliability of the reconstructed bundles. Second, we evaluated the ability of features extracted from the bundles to predict individual differences in cognitive performance. For all analyses, we compared three different methods to reconstruct ODFs – GQI, SS3T, and CSD.

### 3.1 Reliability Analysis

To assess the reliability of reconstructed bundles, we evaluated the fraction of scans for which each bundle could be successfully reconstructed, compared within- and between-subject Dice scores, calculated the discriminability scores for all bundles, compared the bundle reconstructions to the atlas bundles they were based on and calculated the ICC for features extracted from the reconstructed bundles.

#### 3.1.1 Nearly all WM bundles can be reconstructed from single-shell 32- direction DWI scans

Before assessing the reliability of reconstructed bundles across two scans, we first evaluated an even more basic measure: how often a given WM bundle could be reconstructed considering all scans from all subjects. To do this, for each ODF reconstruction method, we calculated the fraction of scans for which a given bundle was produced. For the majority of bundles, reconstruction was successful in nearly all scans across all methods, resulting in a reconstruction success rate close to 1. The mean reconstruction success rates across all bundles were 0.977 for GQI, 0.995 for CSD, and 0.996 for SS3T. There was no numeric difference between methods for most bundles (**Supplementary Fig. S1**). However, for 6 out of 60 bundles, reconstruction success rates were lower for at least one of the three reconstruction methods. These bundles included the left and right optic radiation bundle, the left and right corticobulbar tract, and the dentatorubrothalamic tracts (DRTT) (left-to-right and right-to-left). For these six bundles, the lowest success rates were observed for GQI-based ODFs, whereas using CSD- and SS3T-based ODFs consistently yielded higher success rates. These results establish that most major WM bundles can be reconstructed from single-shell acquisitions even with limited angular resolution.

#### 3.1.2 Reconstructed bundles are reliable across all ODF reconstruction methods

To assess the reliability of the reconstructed bundles, we calculated the Dice scores, reflecting the overlap between two reconstructions of the same bundle (where a score of 0 indicates no overlap and a score of 1 indicates perfect overlap). A bundle with high reconstruction reliability is expected to have a higher Dice score between two scans of the same subject (i.e., within-subject similarity), compared to any two scans of different subjects (i.e., across-subject similarity). We repeated this comparison of within- and between-subject similarity across all bundles and ODF reconstruction methods. **Figure 3A** shows an illustrative example of the left corticospinal tract reconstructed using SS3T-derived ODFs. Across all 60 bundles and all three reconstruction methods, the median within-subject Dice scores were consistently higher than the median between-subject Dice scores (**Figure 3B**; see **Supplementary Fig. S2** for full distributions from all bundles). Nonetheless, there was some variation among methods, with CSD and SS3T having numerically higher Dice scores than GQI. The higher average within-subject Dice score compared to the average between-subject Dice score indicates that bundle reconstructions were reliable and distinguishable across subjects.

**Figure 3:**
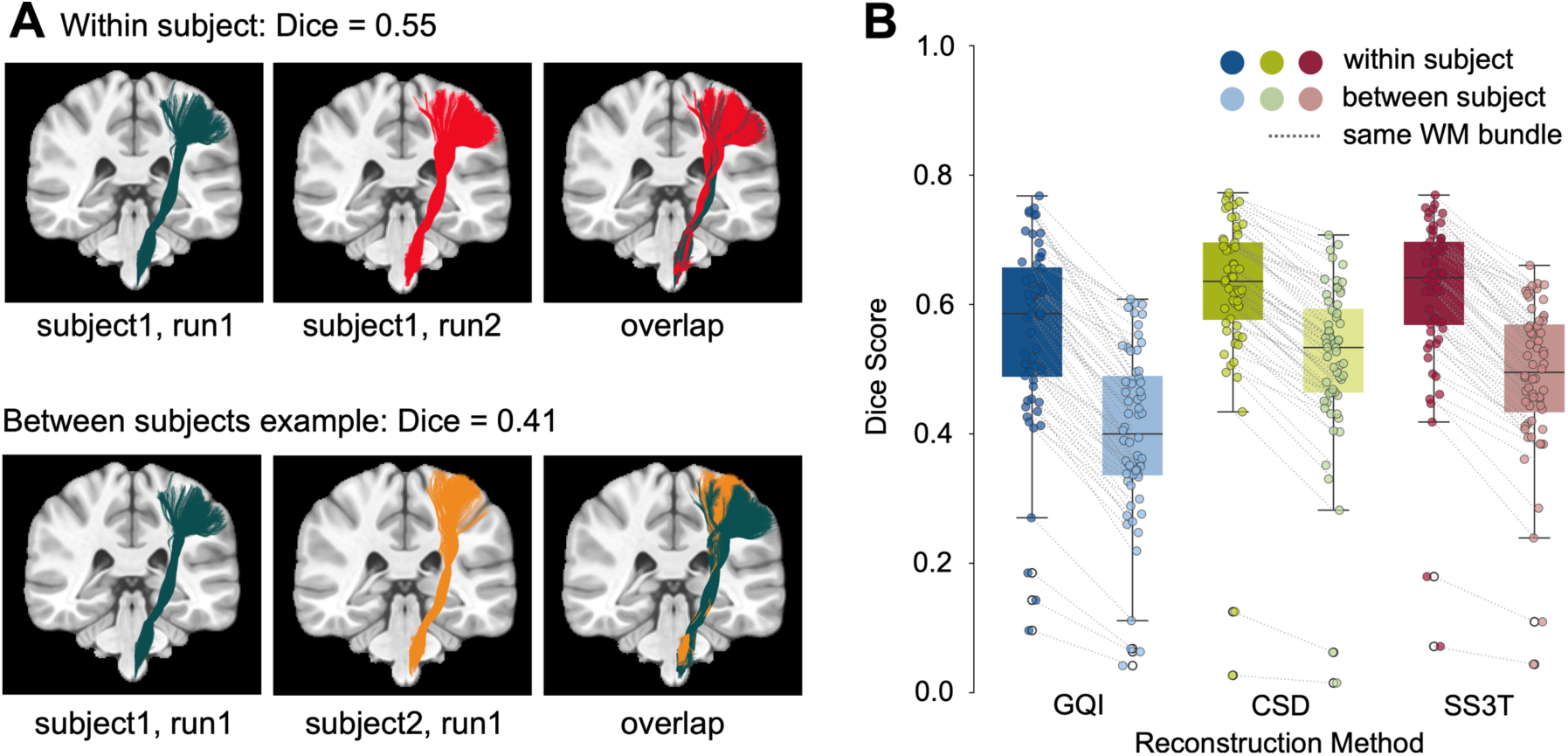
All 60 WM bundles could be reliably reconstructed for all three reconstruction methods. **A)** Example of reconstructed left corticospinal tracts using SS3T-derived ODFs illustrating the higher alignment and corresponding Dice score within subject (first row) compared to between subject (second row). The first row shows the bundle reconstruction for two scans of the same subject, first separately (left and middle) and then overlapping (right). The second row shows an example of scans from two different subjects. Streamline visualizations were created using *MI-Brain* (Rheault et al., 2016). **B)** Median Dice scores within-subject vs. between-subject for each of the three ODF reconstruction methods. Each box plot contains one datapoint per bundle. Within methods, datapoints belonging to the same WM bundle are connected by a dashed line.

#### 3.1.3 Bundles from SS3T have the highest discriminability

To summarize the reliability of the reconstructed WM bundles in one descriptive metric that can be easily compared between reconstruction methods, we calculated the discriminability score for each bundle and each reconstruction method based on the Dice scores. Discriminability is a recently developed measure of test-retest reliability that quantifies how well within-subject observations can be distinguished from between-subject observations. It does so by quantifying the proportion of times that the within-subject distance is smaller than the between-subject distance. All methods had a high median discriminability value across bundles (> 0.94), with SS3T significantly outperforming GQI and CSD (**Figure 4A**). Two outliers were consistently observed across all three methods: the left-to-right and right-to-left DRTTs had the lowest discriminability scores. Notably, these bundles also had the lowest reconstruction success rates. Thus, these bundles were not only challenging to reconstruct but also demonstrated low reliability when reconstructions were available for both scans of a subject.

**Figure 4:**
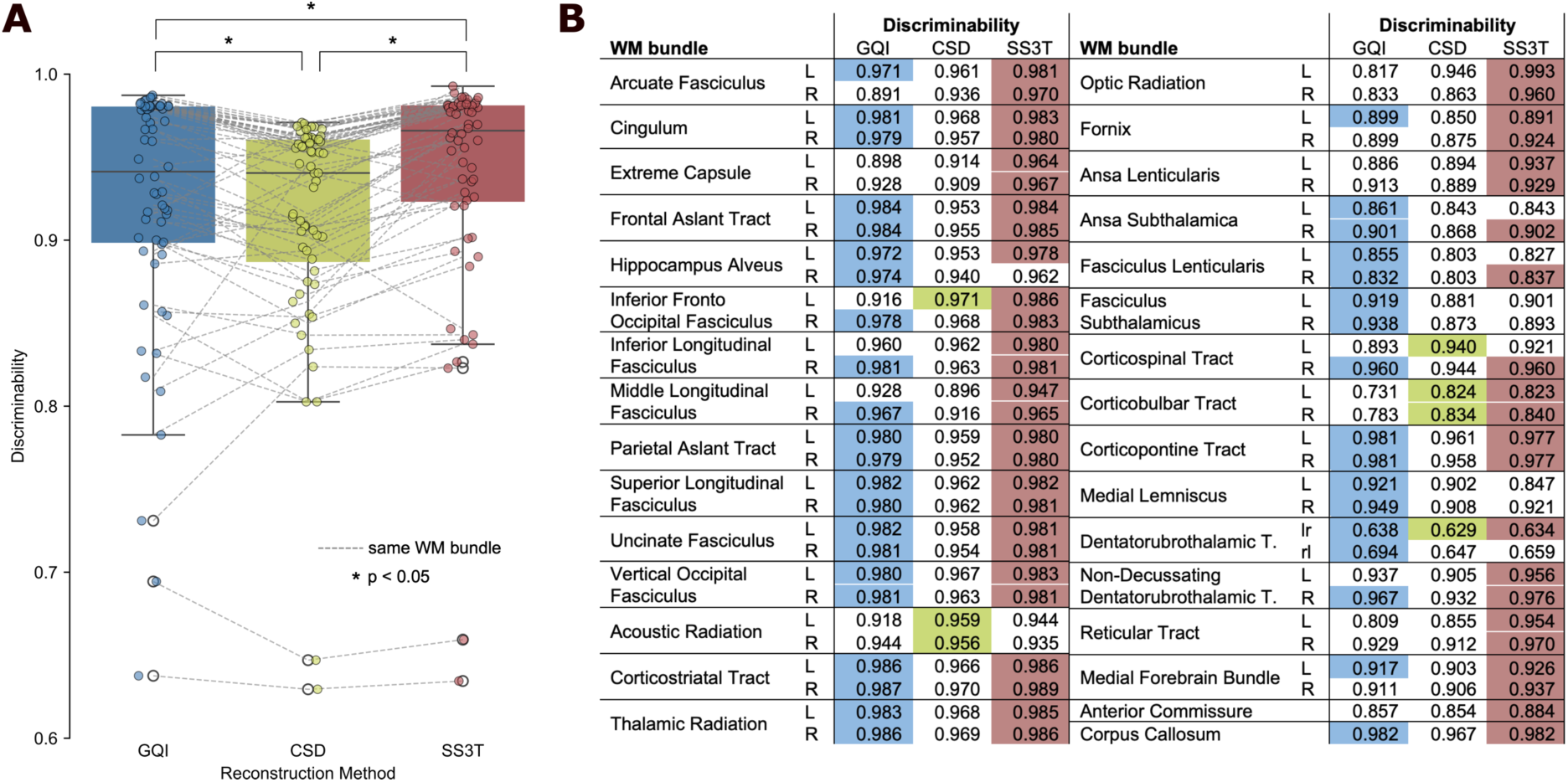
The reconstructed WM bundles exhibit high median discriminability with the highest values for bundles reconstructed from SS3T ODFs. **A)** Plot of the discriminability values for all WM bundles and ODF reconstruction methods. Each data point represents the discriminability value for a specific WM bundle. Corresponding WM bundles are connected with dashed lines across methods. On average, SS3T shows the highest discriminability across reconstruction methods. Significant differences between discriminability distributions (as determined using a Wilcoxon signed rank test and the Benjamini-Hochberg procedure for correction) are marked with an asterisk. **B)** Table showing the exact discriminability values for each of the WM bundles for all three methods, indicating significant differences between methods. A single-colored cell symbolizes that the corresponding method led to the best result and was significantly better than the other methods. Two or three colored cells per bundle show that there was no significant difference between the best two or three methods. Significant differences are determined using permutation tests implemented in *hyppo* (Panda et al., 2020).

As a next step, we statistically compared the discriminability of the three ODF reconstruction methods for each bundle. We found that SS3T was the top-performing method (or tied for top- performing) for 49/60 WM tracts. For GQI, this was true for 39/60 bundles. In contrast, CSD was top (or tied for top) for only 7/60 WM bundles (**Figure 4B**; exact *p*-values can be found in **Supplementary Table S1**). These results suggest that SS3T provides the most discriminable bundle reconstructions, offering the clearest distinction between bundles within vs. across subjects.

#### 3.1.4 Evidence for sensitivity-specificity trade-offs in bundle reconstruction methods

While discriminability indicates whether a bundle is more similar within a subject than between subjects, it does not assess the biological plausibility of the reconstruction. A bundle reconstruction may be highly reliable yet fail to capture all its components, consistently reconstructing only the core of the bundle while omitting peripheral branches. Conversely, a reconstructed bundle could be over-inclusive and consistently include parts of the brain that are not part of the bundle. To evaluate this possibility, we compared the reconstructed bundles to their corresponding atlas bundles (F.-C. Yeh et al., 2018). As described below, we evaluated bundles visually and also calculated sensitivity and specificity values.

For the majority of WM bundles, we observed the following pattern: GQI produced bundles with the highest sensitivity but the lowest specificity, bundles reconstructed using CSD had the highest specificity and lowest sensitivity, whereas SS3T led to a trade-off between sensitivity and specificity (**Supplementary Fig. S3**, **Table S2**). We illustrate this pattern in detail for the left cingulum bundle and the left corticospinal tract.

For the left cingulum bundle, fibers extending to the frontal cortex were missing in all GQI reconstructions, while CSD and SS3T were able to recover these connections more effectively (**Figure 5A**, left). When considering sensitivity and specificity values for the different reconstruction methods, GQI had the lowest sensitivity and the highest specificity – it tended to reconstruct only the core of the bundle, producing few false positives. In contrast, CSD achieved the highest sensitivity but also had the lowest specificity. As such, CSD produced complete reconstructions that often extended over the defined atlas bundles. Finally, SS3T effectively balanced sensitivity and specificity, leading to a more complete reconstruction than GQI with fewer nonspecific connections than CSD (**Figure 5A**, right).

**Figure 5:**
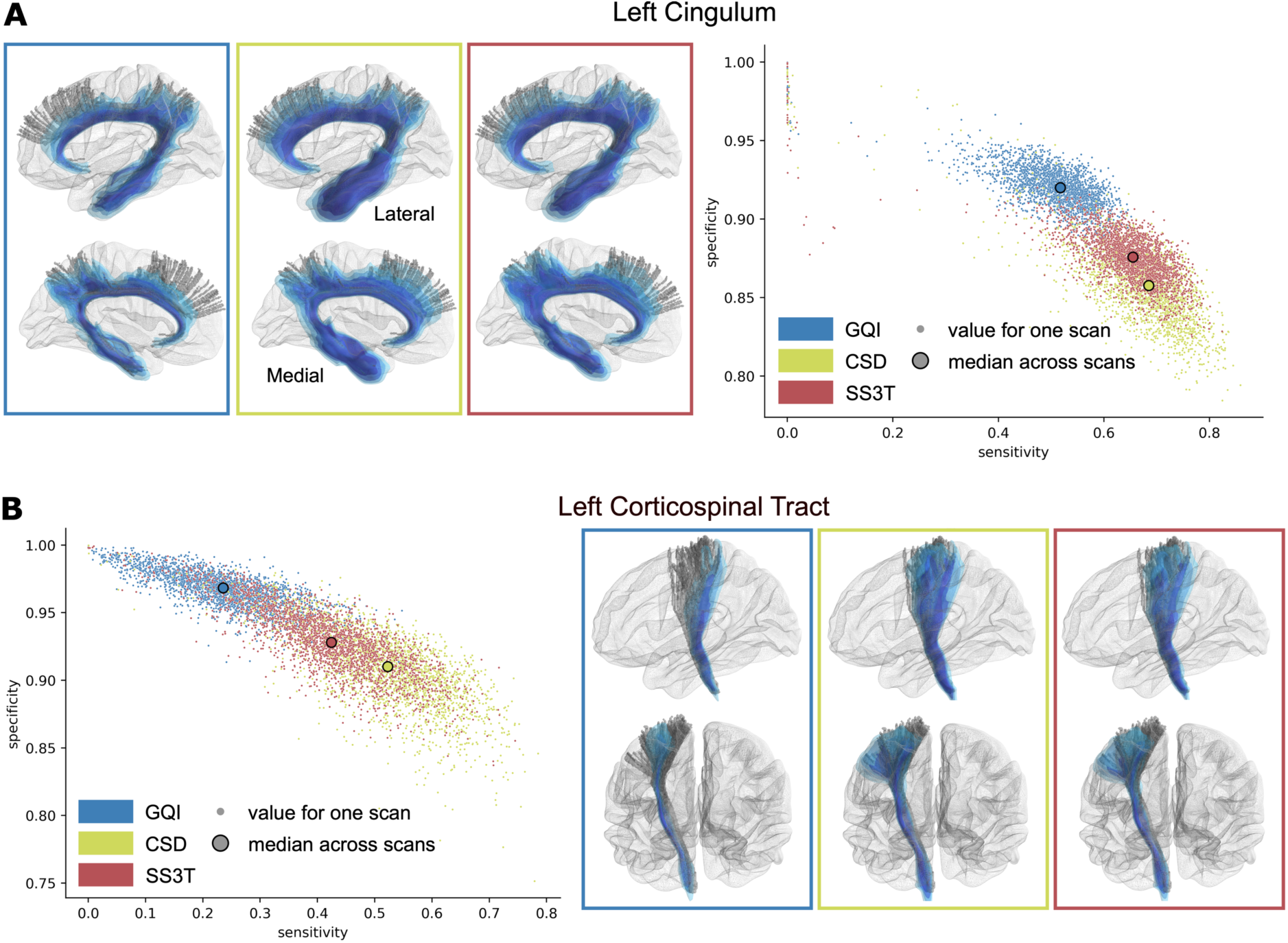
Different ODF reconstruction methods exhibit distinct sensitivity vs. specificity profiles of reconstructed WM bundles when compared to the atlas tracts used for bundle reconstruction. **A)** Left: Example images for the left Cingulum from the lateral and medial views of the left hemisphere. For each reconstruction method, the atlas tract (gray) is overlaid by the probabilistic population map (blues) derived by averaging all reconstructions of the given bundle. Darker colors indicate that the given part of the bundle was reconstructed in a higher fraction of subjects. These plots were created using *Mayavi* (Ramachandran & Varoquaux, 2011). Right: Sensitivity and specificity of each individual reconstructed left Cingulum bundle compared to the corresponding atlas tract (small points). The large points show the median sensitivity and specificity across all scans per method. **B)** Left: Sensitivity and specificity of each individual reconstructed left Corticospinal Tract compared to the corresponding atlas tract (small points). The large points show the median sensitivity and specificity across all scans per method. Right: Example images for the left Corticospinal tract from the lateral and posterior view of the left hemisphere.

We observed a similar pattern for the left corticospinal tract (**Figure 5B**). As for the cingulum bundle, GQI only recovered the core of the bundle, whereas CSD and SS3T reconstructed more of the fibers that branched towards the cortex (**Figure 5B**, left). Again, we found similar sensitivity and specificity patterns for each method as seen for the cingulum bundle (**Figure 5B**, right).

Taken in the context of our prior findings on discriminability, these results emphasize that GQI has relatively high specificity but low sensitivity – it produces discriminable reconstructions by only capturing the core of the bundles. In contrast, CSD is more sensitive but less specific: it provides greater coverage of the atlas bundles but at the cost of reduced discriminability. Finally, SS3T appears to provide a good balance between sensitivity and specificity, providing discriminable and relatively complete reconstructions for most WM bundles.

#### 3.1.5 Features from bundles reconstructed using SS3T are most reliable

The prior steps of the reliability analysis focused on the bundle shape, location, and extent. However, the reconstructed bundles are typically not directly used as input for brain-behavior studies predicting inter-individual differences. Instead, scalar features extracted from the bundles are more commonly used for prediction. Our next step was therefore to assess the reliability of scalar features from WM bundles using intraclass correlation (ICC). Across bundles, ICC values ranged from poor to good depending on the feature and reconstruction method. On average, ICCs were in the fair range (**Figure 6**, exact values per bundle in **Supplementary Table S3**). Further, for all three feature types, SS3T consistently resulted in the significantly highest ICCs across bundles (**Figure 6**). While differences between GQI and CSD were non-significant for FA (**Figure 6B**) and MD (**Figure 6C**), GQI yielded significantly higher ICCs than CSD for bundle volume (**Figure 6A**). The feature type / reconstruction method combination leading to the highest average ICCs across bundles was bundle volume / SS3T. These results highlight that not only the reconstruction method but also the feature type influences reliability.

**Figure 6:**
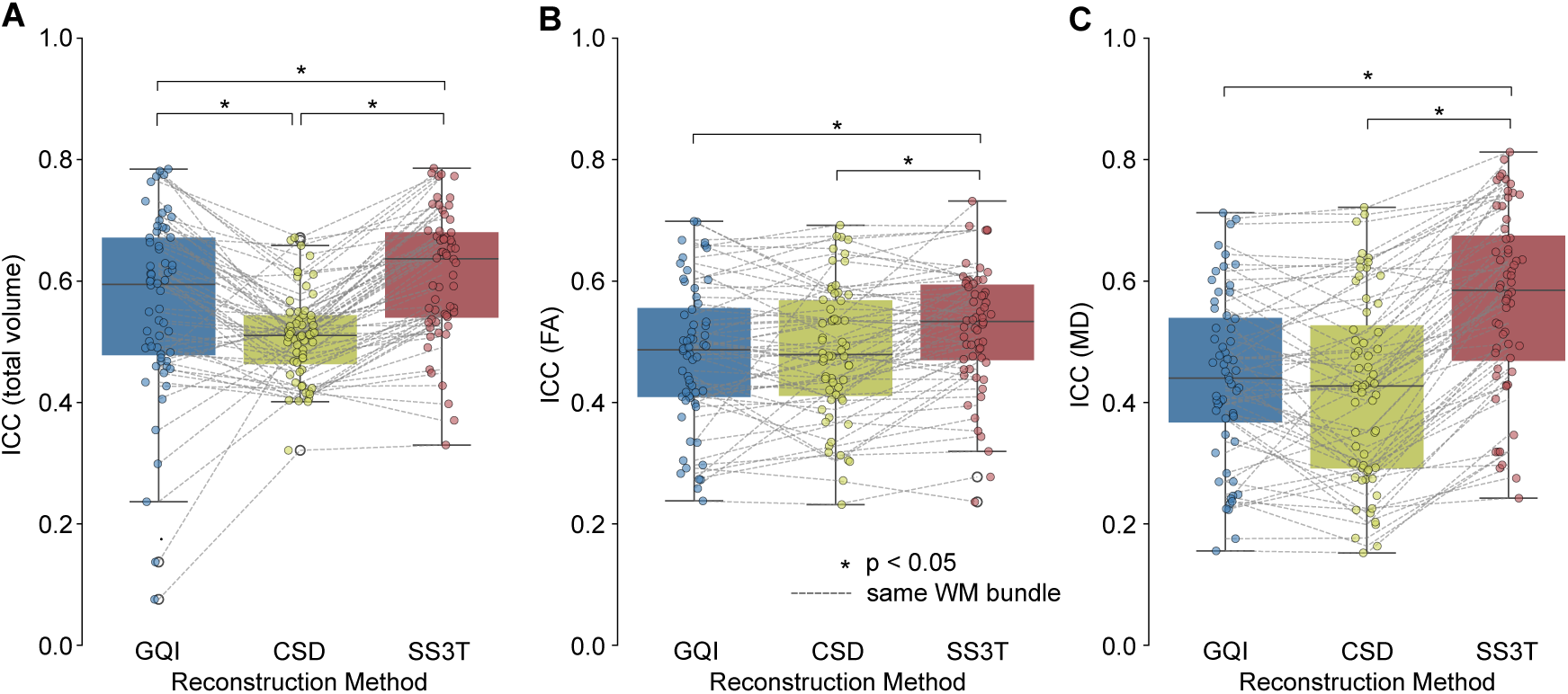
Scalar bundle features (total volume, FA, MD) are most reliable in bundles reconstructed using the SS3T method. ICC was evaluated for **A)** bundle volume, **B)** mean FA across all voxels belonging to the reconstructed bundle, and **C)** mean MD across all voxels belonging to the reconstructed bundle. Each data point represents the ICC value for a specific WM bundle. Corresponding WM bundles are connected with dashed lines across methods. Significant differences after correction for multiple comparisons are marked with an asterisk.

### 3.2 Prediction Analysis

As a next step, we aimed to assess how well features extracted from single-shell scans with limited angular resolution are suited for predictive analyses that link brain and behavior. Specifically, we evaluated how well volumetric and microstructural features from WM bundles could predict individual differences in complex reasoning performance. As described below, we compared the prediction accuracy and prediction similarity between the three ODF reconstruction methods for four different feature groups.

#### 3.2.1 GQI and SS3T outperform CSD for predicting complex reasoning

We considered a total of 24 prediction frameworks (4 feature groups x 3 reconstruction methods x 2 scans) for predicting complex reasoning. Across configurations, the median test Pearson correlation between the ground truth and predicted cognition scores, reflecting the prediction accuracy, ranged from *r* = 0.15 to *r* = 0.36 (**Figure 7**). When comparing the different ODF reconstruction methods, features from bundles based on GQI and SS3T reconstruction outperformed features from bundles based on CSD for prediction in 15/16 comparisons. Although the differences reached statistical significance for 5 out of 16 comparisons, the overall pattern consistently showed GQI and SS3T outperforming CSD. Notably, there was no consistent performance difference between SS3T and GQI (**Figure 7**). When comparing the influence of bundle feature types, we found that using all features led to the best prediction of complex reasoning. When examining bundle features separately, prediction accuracy was best for volume, followed by FA, and finally MD. This was consistent across both scans and all three reconstruction methods (**Figure 7**). These patterns were consistent when considering the MSE as a proxy for prediction accuracy (**Supplementary Fig. S4**). Notably, these findings parallel our reliability results and suggest that methods that more reliably reconstructed bundles, like SS3T and GQI, allow for better prediction performance of cognition.

**Figure 7:**
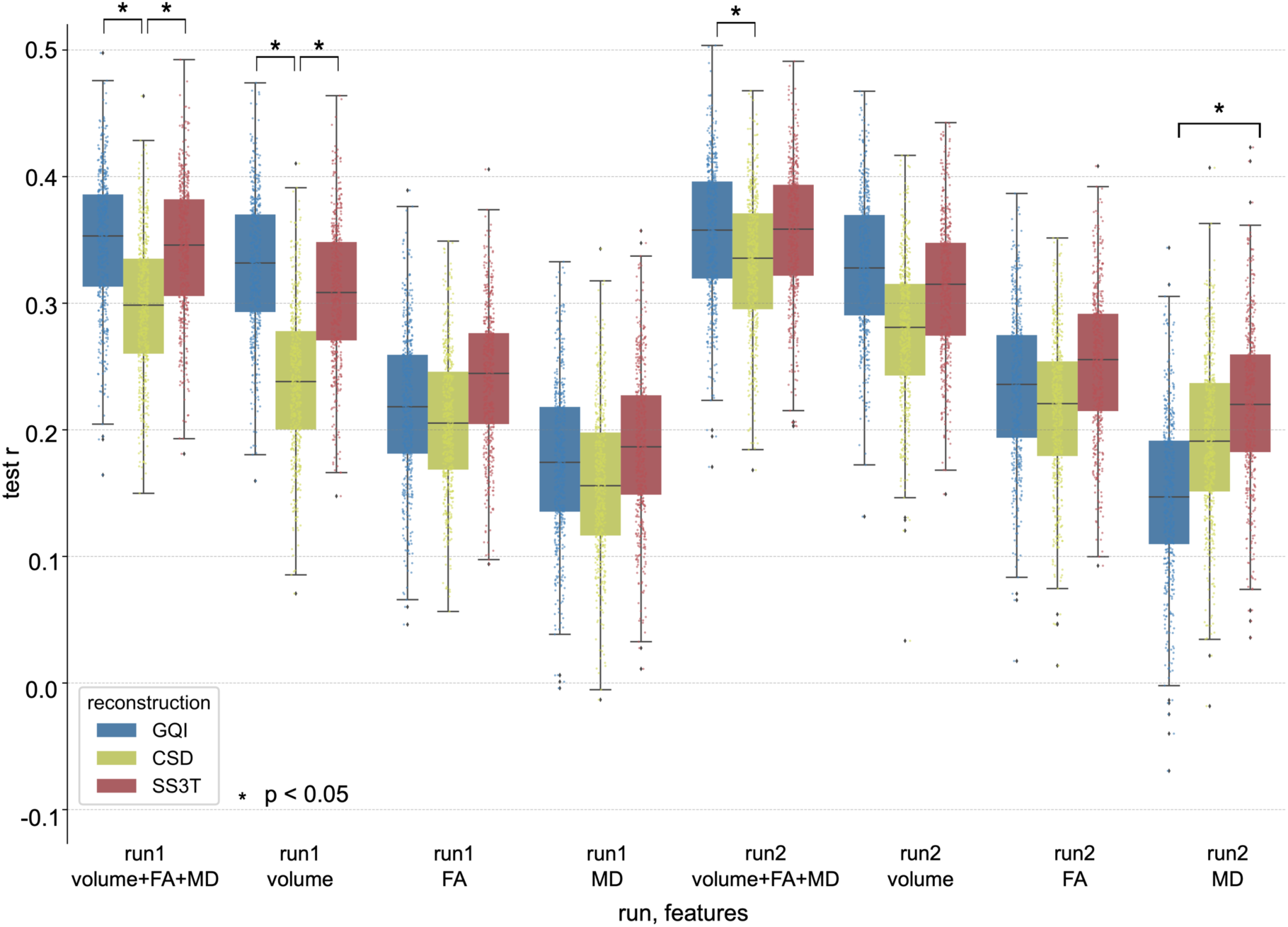
GQI and SS3T outperform CSD in predicting complex reasoning from different groups of bundle features. Each distribution contains 500 points (100 x 5-fold CV). The left block of prediction accuracies (Pearson r) used features extracted from bundles reconstructed from run-01 scans, and the right block from run-02 scans. For each run, four different groups of features were evaluated: volume, mean FA, and mean MD for each of the 54 considered bundles (162 features), only the bundle volume (54 features), only the mean FA (54 features,) and only the mean MD (54 features). Significant differences between the different reconstruction methods, within run and feature groups, are marked with an asterisk. Significance was determined with an adjusted t-test implemented in *Julearn* (Hamdan et al., 2023) and corrected for multiple comparisons using the Benjamini-Hochberg procedure.

#### 3.2.2 Predictions remain similar when using features from two different scans for prediction

To assess how consistent the predictive models were across scans, we compared the predictions generated from features derived from the first and second scans separately. Specifically, we used the same model and cross-validation splits to generate predictions from each scan and then calculated the Pearson correlation between these two sets of predicted scores. Similar to the findings for prediction accuracy and feature ICC, models using GQI and SS3T features produced more similar predictions across runs than those using CSD features (**Figure 8**). The difference between the methods was strongest when using only bundle volume as a feature. Overall prediction similarity was highest when considering all features or only the bundle volumes as features. (**Figure 8**). These results further emphasize the utility of SS3T and GQI for brain-behavior prediction analyses and stress the importance of selecting a reliable reconstruction method.

**Figure 8:**
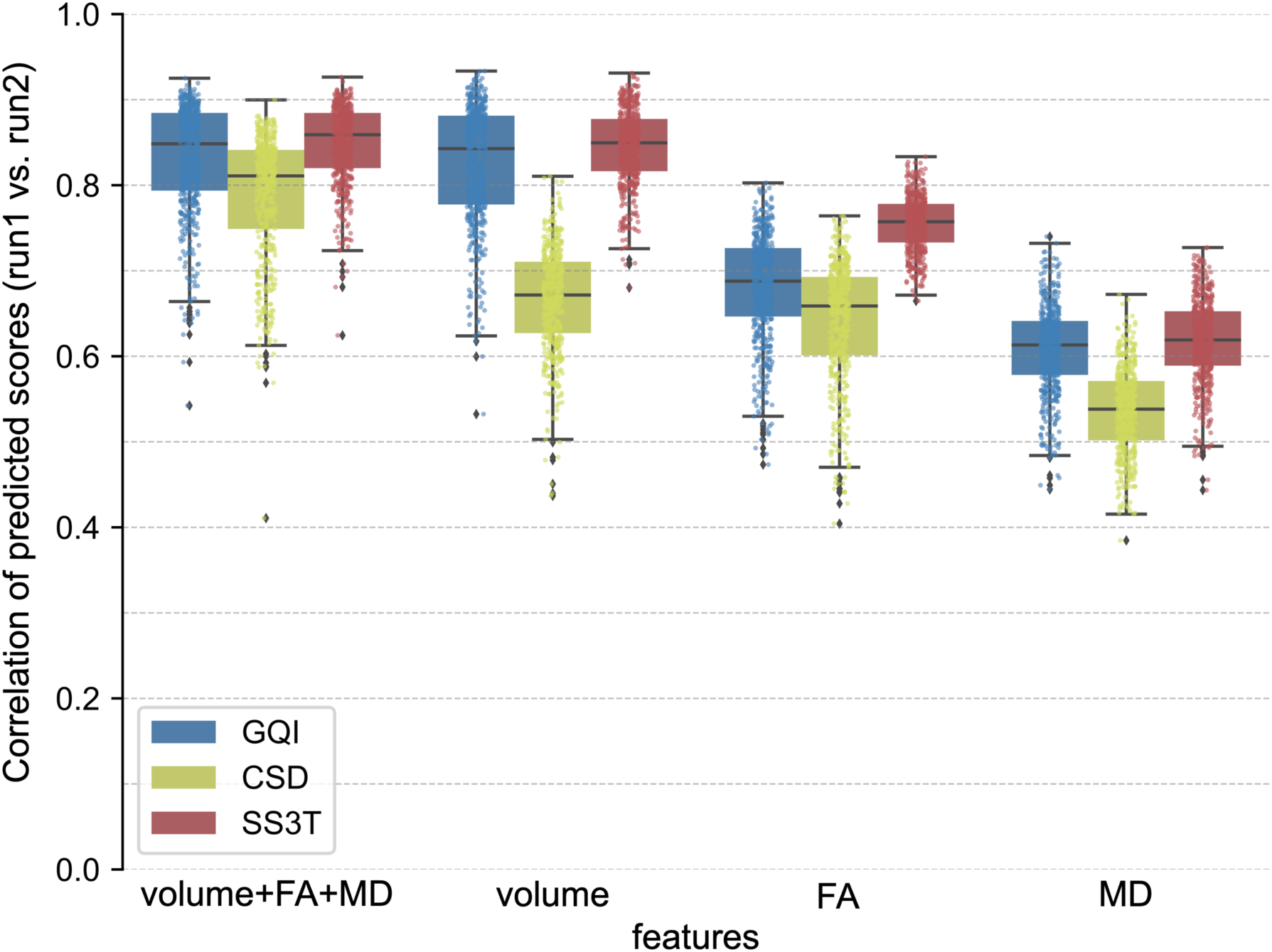
Predictions of complex reasoning remain similar when using features from two different scans for prediction, with GQI and SS3T outperforming CSD. Comparison of the prediction similarity between different reconstruction methods for various groups of features. Prediction similarity was assessed by correlating the predicted scores obtained from features extracted from run-01 scans with predictions based on run-02 features for each fold. A higher correlation indicates a higher similarity across scans. Each distribution contains 500 points (100 x 5-fold CV).

#### 3.2.3 Prediction results are similar across cognitive domains

To test the robustness of our findings, we conducted two sensitivity analyses. First, we assessed whether prediction results were consistent across different cognitive scores: overall accuracy and executive functioning. Second, we evaluated the impact of adjusting for total brain volume (TBV) in our analyses.

When predicting the overall accuracy, median prediction accuracy was in the range *r* **∈** [0.06, 0.28] (**Supplementary Fig. S5A**). For executive functioning, correlations were in the range of *r* **∈** [0.10, 0.21] (**Supplementary Fig. S5B**). As for the main analysis, GQI and SS3T outperformed CSD. Similarly, all features performed better than only volume, only FA, and only MD for both prediction accuracy (**Supplementary Fig. S5**) and prediction similarity (**Supplementary Fig. S6**). For the similarity of prediction results between two DWI scans, as in the main analysis, the difference was largest when considering only the bundle volumes as features (**Supplementary Fig. S6**). These results confirm that the patterns observed in the main prediction task generalize to other cognitive scores.

When controlling for TBV when predicting complex reasoning, we observed an overall drop in prediction accuracies to *r* **∈** [0.06, 0.18] (**Supplementary Fig. S7A**). While SS3T and GQI continued to outperform CSD for prediction accuracy (**Supplementary Fig. S7A**) and prediction similarity (**Supplementary Fig. S7B**), the difference between feature groups decreased. Combining all three feature types still led to improved performance but with less difference between different feature types (**Supplementary Fig. S7**). These results show that the ranking of reconstruction methods persists when including TBV as a covariate. However, it also suggests that the predictive power of bundle volume features may be partly driven by the global effects of TBV.

## 4 Discussion

Our findings demonstrate that single-shell DWI scans with lower angular resolution can serve as a valuable resource for research. While certain white matter bundles posed challenges in reconstruction, the majority were successfully reconstructed and exhibited high reliability across repeated scans. By providing precise values for discriminability, sensitivity, specificity, and ICCs for each bundle and reconstruction method, our results offer researchers benchmark data to help select reconstruction strategies tailored to their specific research questions. Importantly, we showed that these reliably reconstructed bundles were not only reproducible but also informative for brain-behavior analyses predicting cognitive performance. Notably, as the scans evaluated here are quite similar to those acquired as part of both legacy research studies and clinical practice at some academic centers (e.g. the Children’s Hospital of Philadelphia; (Zimmerman et al., 2025), these results underscore the substantial research potential of single-shell scans with lower angular resolution.

In tractography analysis using dMRI, it is well-established that different processing pipelines can lead to different outcomes (Brun et al., 2019; Neher et al., 2015; Schilling et al., 2019). This prior work motivated our evaluation of three ODF reconstruction methods. We included CSD based on its ubiquity and its compatibility (after file format conversion) with *AutoTrack*, which we used for tract segmentation. GQI served as a standard ODF reconstruction method commonly paired with *AutoTrack*. Finally, SS3T was included as it has been optimized for single-shell data. Notably, we did not include diffusion tensor imaging (DTI) (Le Bihan et al., 2001), as it can only reconstruct one major diffusion direction per voxel and shows poor tractography performance (Berman et al., 2013; Kamagata et al., 2024; Petersen et al., 2017). Overall, our findings highlight an advantage of using SS3T for ODF reconstruction on single-shell data with 32 diffusion directions, as it leads to both complete and reliable bundle reconstructions that perform well for cognitive prediction. However, depending on the tract of interest and the goals of the study – e.g., prioritizing coverage or discriminability – alternative methods such as CSD or GQI may be more appropriate in specific contexts. For example, if sensitivity is more important than specificity or reliability, CSD should be considered. In contrast, when wanting to accurately extract microstructural properties from the core of a bundle while minimizing the impact of false positives on the measurements, GQI might be the best choice considering its high specificity.

While most prior comparisons of tractography pipelines have focused on dMRI data of different qualities (Baete et al., 2019; Daducci et al., 2014), one in-depth study evaluated multiple ODF reconstruction methods on lower angular resolution data with a *b*-value of 1000 s/mm² (Wilkins et al., 2015). This study identified CSD and the ball-and-stick model as yielding the best fiber detection rates (Wilkins et al., 2015). Although SS3T was not available at the time, GQI was included and showed lower coverage but fewer false positives, aligning with our current findings. That study, however, used only a single scan per subject and, as such, could not assess test-retest reliability. In contrast, our results show that while CSD consistently produces more complete bundles, this does not necessarily translate into greater reproducibility across repeated scans.

Regardless of the reconstruction method, most bundles could be reliably identified from these limited scans, as reflected in generally high rates of successful reconstruction. This is in line with what would be expected from advanced acquisitions (F.-C. Yeh, 2022). The DRTT was a notable exception, exhibiting consistently low reconstruction success across methods. The DRTT is a long tract connecting the dentate nucleus in the cerebellum to the motor cortex via the red nucleus and thalamus. Prior work suggests that it is difficult to reconstruct due to its multisynaptic nature and interhemispheric course (Kwon et al., 2011). Its reconstruction is known to be challenging even in advanced diffusion MRI datasets (Radwan et al., 2022), with reported reconstruction success rates of 0.625 in the Human Connectome Project dataset (Van Essen et al., 2013) and complete failure in the MASSIVE dataset (Froeling et al., 2017). These prior reports underscore that limitations in DRTT reconstruction were not solely attributable to the lower angular resolution, single-shell data used in our analysis.

Beyond the DRTT, our results indicate that WM bundles can be reconstructed reliably despite the limits of the acquisition sequence. Mean within-subject Dice scores were consistently higher than between-subject scores across all bundles and reconstruction methods, though absolute within-subject values remained moderate (averaging around 0.6). Prior work has shown that removing two different directions from a single 100-direction scan leads to Dice scores around 0.87 (Vos et al., 2016), stressing how even minor acquisition differences reduce reliability. In high-quality diffusion spectrum imaging (DSI) data with 258 directions, median within-subject Dice scores around 0.80 have been reported across WM bundles (Radhakrishnan et al., 2023). Here, when considering only the bundles also included in (Radhakrishnan et al., 2023), median within-subject Dice scores of 0.59 (GQI), 0.65 (CSD), and 0.65 (SS3T) were observed. This showed that while reliable bundles could be reconstructed from limited scans, reliability is lower than cutting-edge research scans.

It is worth emphasizing that comparing dice values across the literature can be challenging and requires nuance. Many prior studies have reported the weighted Dice score (wDice). This metric weights voxels by their streamline density when assessing overlap, giving more weight to areas with dense fibers, such as the core of the bundle, and less weight to spurious streamlines that are far from the core of the fascicle, e.g., branching out at the cortex (Cousineau et al., 2017). The wDice score has been used frequently for evaluating the within-subject similarity of reconstructed WM bundles (Boukadi et al., 2019; Cousineau et al., 2017; Kruper et al., 2021; Radwan et al., 2022; F. Zhang et al., 2019). It is, however, difficult to compare it to the unweighted Dice score we used here. We deliberately decided against using the wDice score because it has been shown that the streamline density often does not reflect underlying biology but rather how difficult it is to track through a considered voxel (C.-H. Yeh et al., 2021) based on regions of crossing fibers or a bottleneck effect (Schilling et al., 2022). Further, it may not be optimal for our purpose of comparing both within- and between-subject overlap. Since wDice places more emphasis on densely tracked regions, which tend to be more similar across subjects, it reduces sensitivity to inter-subject differences and may thus inflate estimates of reliability.

In addition to the Dice score, we used discriminability to summarize the reliability per bundle. We found that WM bundles from low-resolution scans can be identified with high accuracy, especially when reconstructed using SS3T, which outperformed GQI and CSD. Discriminability values varied between 0.629 and 0.993 for different bundles and ODF reconstructions, with means >0.94. While there is no comparable work evaluating the discriminability of reconstructed WM bundles, studies examining the reliability of functional connectivity have reported discriminability values ranging from 0.654-0.98 (Bridgeford et al., 2021; Camp et al., 2024; Shearer et al., 2025). For inter-regional structural connectivity in research quality data, one study reported that discriminability varies by the number of parcels in the GM atlas and with the type of edge weighting, with values between 0.75 and 0.96 (Bridgeford et al., 2021). As such, our results for single-shell scans with limited angular resolution are well within the range of what could be expected for widely used neuroimaging derivatives.

While a reliable bundle reconstruction is essential, most brain-behavior studies leverage scalar features derived from WM bundles, such as total volume, FA, and MD. We found that scalar features extracted from reconstructed bundles showed fair test-retest reliability as defined by the ICC. Prior work has reported that the volume of association bundles reconstructed from the HCP-YA dataset had a median ICC of 0.81 (F.-C. Yeh, 2020). Considering only equivalent bundles, we observed lower but still acceptable median ICCs for bundle volume: 0.63 for GQI, 0.52 for CSD, and 0.67 for SS3T. Importantly, these values remained within the range of other widely used neuroimaging features employed successfully in inter-individual prediction. For example, resting state functional connectivity (rsFC) exhibits ICCs between 0.15 and 0.65 with an overall mean of 0.29 (Noble et al., 2019). For task FC, these values range from -0.02 to 0.87, with a mean of 0.397 (Elliott et al., 2020). For inter-regional structural connectivity, reported ICCs have ranged from 0.35 to 0.62 across reconstruction methods (Buchanan et al., 2014) and 0.50 to 0.67 for different types of data (Prčkovska et al., 2016). As such, features derived from single-shell dMRI with lower angular resolution have ICCs similar to other measures widely used in contemporary research.

In addition to possessing acceptable reliability, the results of our brain-behavior analyses demonstrated that WM bundles reconstructed from limited diffusion acquisitions can predict cognitive performance (mean prediction accuracy *r* between 0.06-0.36). While relatively few studies have used features derived from WM bundles for cognition prediction, existing approaches primarily rely on the advanced diffusion data from the HCP Young Adult (HCP-YA) dataset (Van Essen et al., 2013). When not correcting for any confounding factors, mean prediction accuracies between *r* = 0.053 and 0.335 could be observed for different cognition scores and features (Lo et al., 2025). When TBV was accounted for, prediction accuracies ranged from *r* = 0.18 to 0.37 (W. Liu et al., 2023). Similar results have been observed in studies predicting cognition from the structural connectome (SC) derived from advanced datasets such as the HCP-YA and ABCD datasets (Garavan et al., 2018), with values comparable to ours: *r* = 0.05-0.3 (Z. Zhang et al., 2019), 0.13-0.41 (M. Liu et al., 2021), 0.2-0.3 (Dhamala et al., 2021), and -0.02-0.25 (Rauland et al., 2025). When accounting for confounds such as age, sex, education, and motion, prediction performance was in the range of: *r* = -0.02-0.32 (Y. Zhang et al., 2024), -0.02-0.26 (Litwińczuk et al., 2022), and 0.15-0.28 (Ooi et al., 2022). Despite relying on clinically feasible single-shell, lower angular resolution scans and rigorously controlling for relevant confounds, our prediction results are thus on par with studies that use much more advanced diffusion data. This underscores the potential of such limited dMRI acquisitions for brain-behavior studies.

As part of our brain-behavior analyses, we observed that bundle volume consistently outperformed FA and MD in prediction accuracy, aligning with prior findings (Yeung et al., 2023; Z. Zhang et al., 2019). However, this superior performance might have been partially driven by the global effects of TBV (Pietschnig et al., 2015; Royle et al., 2013). Including TBV as a confound and regressing it from the features resulted in a general drop in prediction performance. Nevertheless, the remaining signal indicates that WM bundle features contain information relevant to cognition beyond TBV itself. Furthermore, combining all three feature types improved prediction performance across reconstruction methods, though the comparison is partially confounded by the difference in feature dimensionality (i.e., 162 vs. 54 features).

Several limitations should be considered when interpreting our results. First, although our data were acquired using clinically feasible parameters, they were collected in a research setting. Actual clinical data may introduce additional sources of variability and artifacts, such as increased motion in patient populations or structural abnormalities like tumors, stroke lesions, or other pathological changes. Moreover, clinical diffusion MRI protocols vary across institutions and countries. While the presented acquisition parameters are feasible to be applied in large academic centers like the Children’s Hospital of Philadelphia, it is not uncommon for hospitals to acquire images with as few as 6 gradient directions, which is the minimum necessary number of directions to model the diffusion tensor. We do not expect the results to generalize to scans with fewer gradient directions. Second, our study was conducted using data from a single site and scanner; when pooling data from multiple sites or over a longer time in which the institution’s scanner, acquisition protocol, or software might have changed, harmonization (Moyer et al., 2020; Pinto et al., 2020) becomes necessary. Third, while our two 32-direction scans were representative of clinical constraints, they were originally designed to be combined into a single 64-direction scan and thus used different sets of gradient directions. Compared to traditional test-retest data, this may have introduced additional variability in the data. However, this limitation would lead to an underestimation rather than an overestimation of reliability. Finally, as in Wilkins et al (2015), we would like to stress that the relatively low *b*-value and the low number of directions might not be equally well suited for all three reconstruction methods. Generally, all three methods evaluated here are optimized for higher *b*-values (GQI *b*=1500, CSD: *b*=3000, SS3T: *b*=2500) and a higher angular resolution (GQI: >100, CSD: >60, SS3T: >45), and their relative performance may change for acquisitions including a higher *b*-value and more directions.

In summary, this study highlights the research potential of single-shell dMRI data with relatively limited angular resolution. We demonstrated that most major white matter bundles can be reliably reconstructed and that features derived from these reconstructions are reliable enough to allow for brain-behavior analyses. In addition, we show that SS3T generally provided the highest discriminability while still reconstructing relatively complete bundles. The results lay the groundwork for future studies to apply modern analytic methods, such as person-specific bundle segmentation, to the massive pool of legacy research dMRI data and clinically acquired dMRI scans. Notably, healthcare systems routinely collect diffusion scans at a scale unmatched by any research consortium, with samples that are more demographically diverse and representative of the general population. Unlocking this resource would not only allow the creation of more generalizable normative brain charts (Schabdach et al., 2023), but also enable studies of rare diseases that are difficult to capture in research cohorts. Further, truly massive dMRI datasets have the potential to both improve brain-behavior analyses (Cui & Gong, 2018; Gell et al., 2024; Marek et al., 2022; Scheinost et al., 2019) and may be critical for leveraging artificial intelligence methods that require massive training datasets (Hoffmann et al., 2022).

## Acknowledgments

This study was supported by grants from the National Institute of Health: R01MH120482 (T.D.S..), R01MH113550 (T.D.S.), R01MH112847 (R.T.S. and T.D.S.), R01MH123550 (R.T.S.), R01NS060910 (R.T.S.), R01MH123563 (R.T.S.), R01MH120174 (D.R.R.), R01MH132934 (A.A.B.), R01MH133843 (A.A.B.), R01MH117014 and R01MH119219 (R.C.G. and R.E.G.), K23MH133188 (E.B.B), T32MH019112 (S.L.M.), T32MH016804 (V.J.S.), BBRF NARSAD #31319 (E.B.B.), F31MH136685 (J.B.), and by the German Research Foundation (Deutsche Forschungsgemeinschaft, DFG) – grant number 269953372/GRK2150. Additional support was provided by the Penn-CHOP Lifespan Brain Institute, the AE Foundation, and the “ML hours” initiative from INM-7, FZJ.

## Data and Code Availability

This paper analyzes existing, publicly available data from the Philadelphia Neurodevelopmental Cohort. Imaging and demographic data are available at the database of Genotypes and Phenotypes (dbGaP): https://www.ncbi.nlm.nih.gov/projects/gap/cgi-bin/study.cgi?study_id=phs000607.v3.p2. To request access, an authorization request can be completed via https://dbgap.ncbi.nlm.nih.gov/aa/wga.cgi?%20page=login&page=login. All original code for image preprocessing, bundle reconstruction, reliability and prediction analysis, as well as code for creating the figures, has been deposited at https://github.com/PennLINC/clinical_dmri_benchmark. A detailed guide to the code and procedures for replicating all analyses can be found at https://pennlinc.github.io/clinical_dmri_benchmark/.

## Author Contributions

**Conceptualization**: M.C., T.D.S., **Methodology**: A.R., S.L.M., A.F.A.-B, M.C., T.D.S, **Software**: A.R., S.L.M., E.B.B., V.J.S., M.C., **Validation**: S.L.M, **Formal analysis**: A.R., **Resources**: S.B.E., T.D.S., **Data Curation**: R.E.G., R.C.G., T.M.M., D.R.R., T.D.S., **Writing - Original Draft**: A.R., **Writing - Review and Editing**: A.R., S.L.M., A.F.A.-B., J.B., E.B.B., R.E.G., R.C.G., A.C.L., T.M.M., O.V.P., K.R., D.R.R., R.T.S., S.S., V.J.S., A.V., S.B.E., M.C., T.D.S., **Visualization**: A.R., M.C., **Supervision**: O.V.P., K.R., S.B.E., M.C., T.D.S., **Funding acquisition**: S.B.E., T.D.S.

## Supplementary

**Figure S1:**
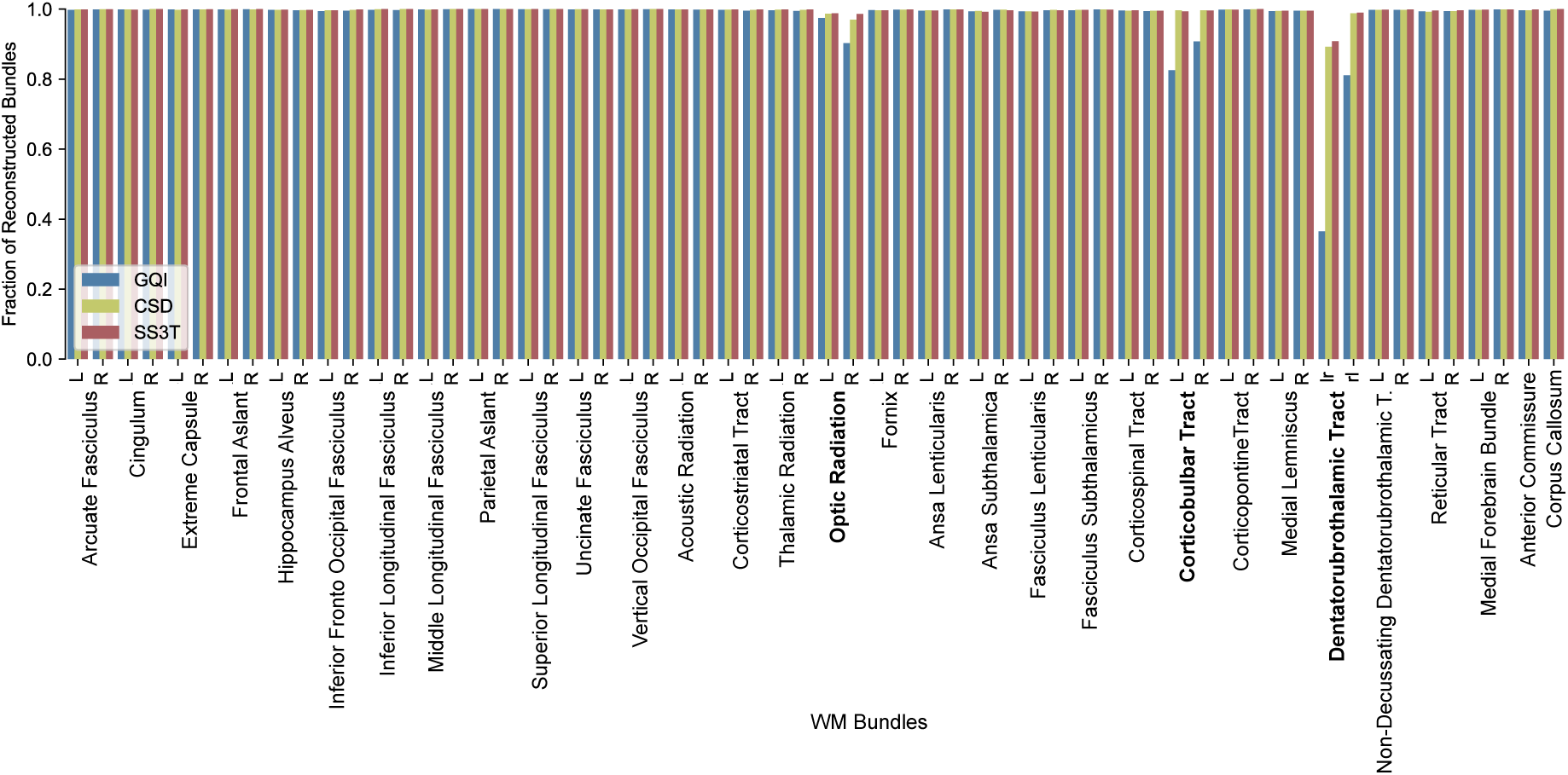
Reconstruction success rates per bundle per method. The reconstruction success rates represent the fraction of scans for which a given bundle could be reconstructed. Marked in bold are bundles with reconstruction fractions < 1 for at least one of the three reconstruction methods.

**Figure S2:**
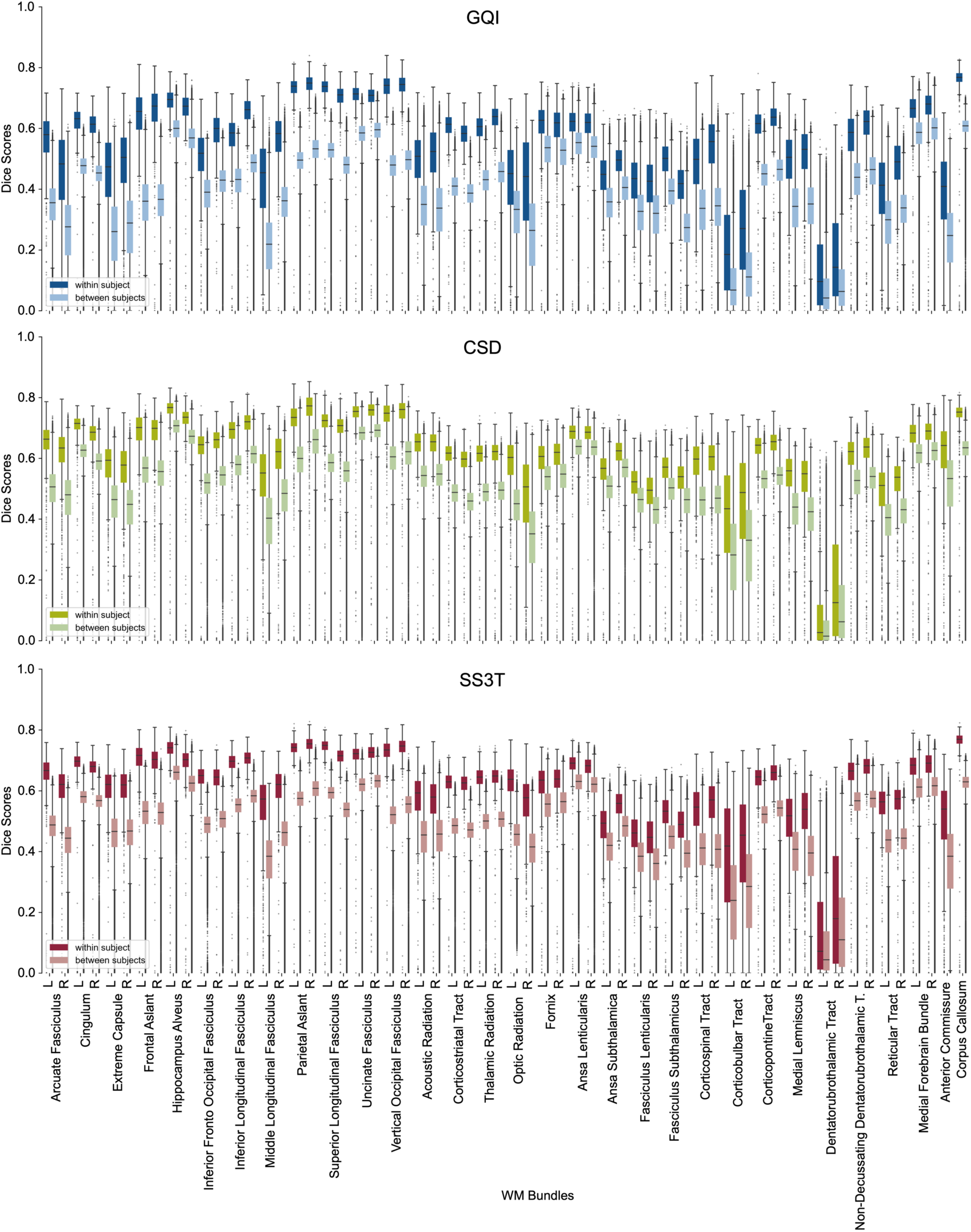
Full distributions of within and between-subject dice scores per bundle for GQI, CSD, and SS3T.

**Figure S3:**
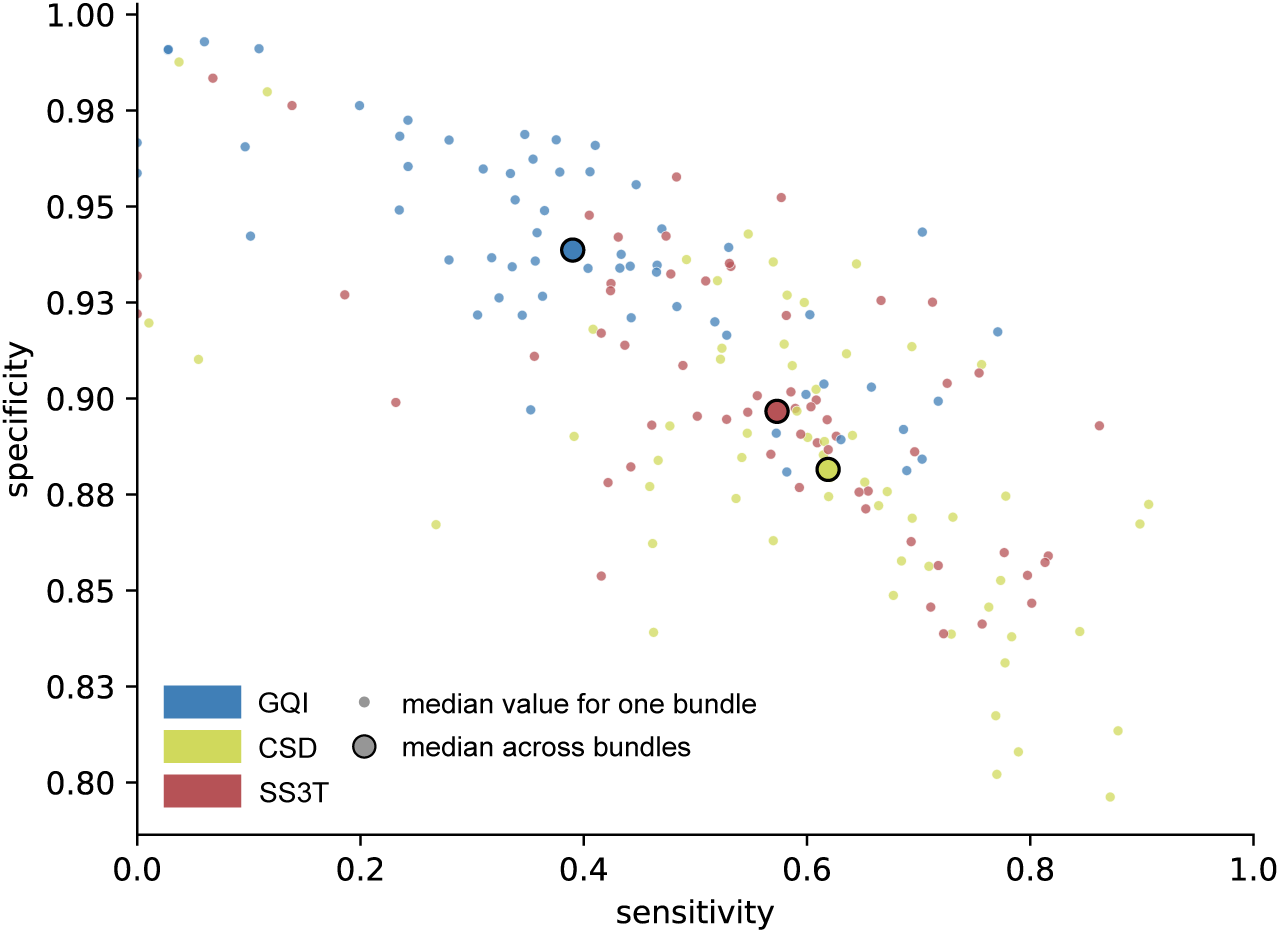
Median sensitivity and specificity for all 60 reconstructed WM bundles. Sensitivity and specificity were calculated for each instance (different subjects, different scans) of the reconstructed bundle. Visualized here are the median sensitivity and specificity values across all instances of a given bundle.

**Figure S4:**
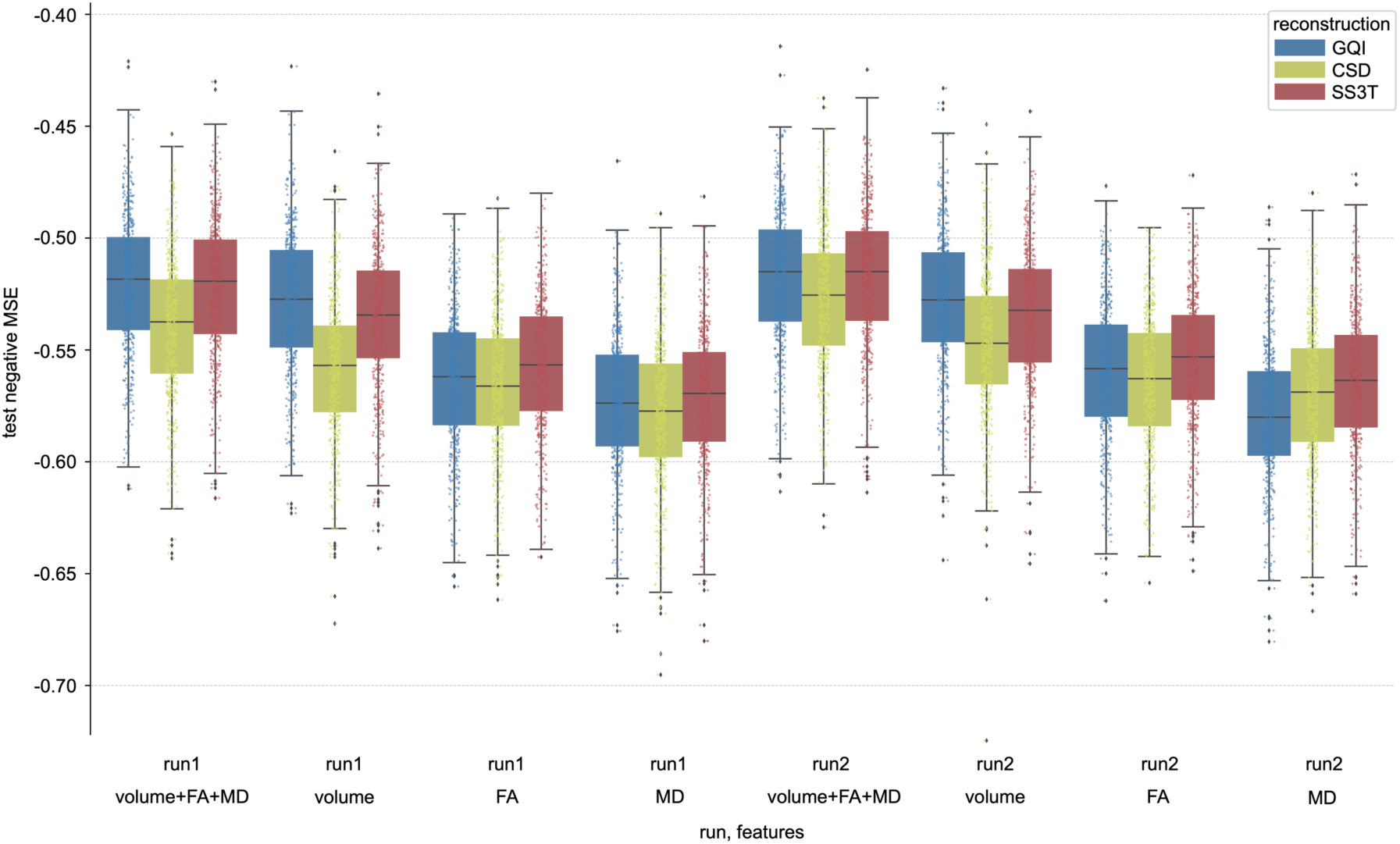
Prediction accuracy in terms of negative MSE for predicting complex reasoning from different groups of features for all three reconstruction methods. Each distribution contains 500 points (100 x 5-fold CV). The left block of prediction accuracies used features extracted from bundles reconstructed from run-01 scans, the right block from run-02 scans. For each run, four different groups of features were evaluated: volume, mean FA and mean MD for each of the 54 considered bundles (162 features), only the bundle volume (54 features), only the mean FA (54 features), and only the mean MD (54 features).

**Figure S5:**
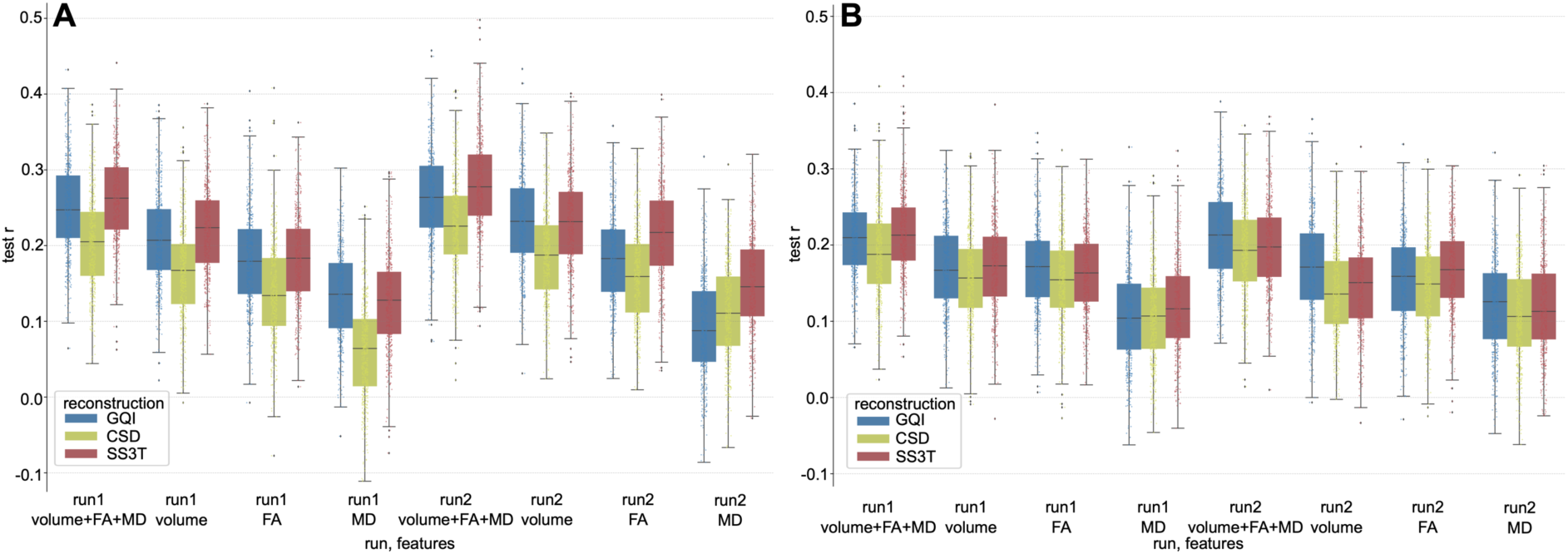
Prediction accuracy in terms of Pearson correlation for predicting **A** a composite IQ score and **B** executive functioning from different groups of features for all three reconstruction methods. Each distribution contains 500 points (100 x 5-fold CV). The left block of prediction accuracies used features extracted from bundles reconstructed from run-01 scans, and the right block from run-02 scans. For each run, four different groups of features were evaluated: volume, mean FA, and mean MD for each of the 54 considered bundles (162 features), only the bundle volume (54 features), only the mean FA (54 features), and only the mean MD (54 features).

**Figure S6:**
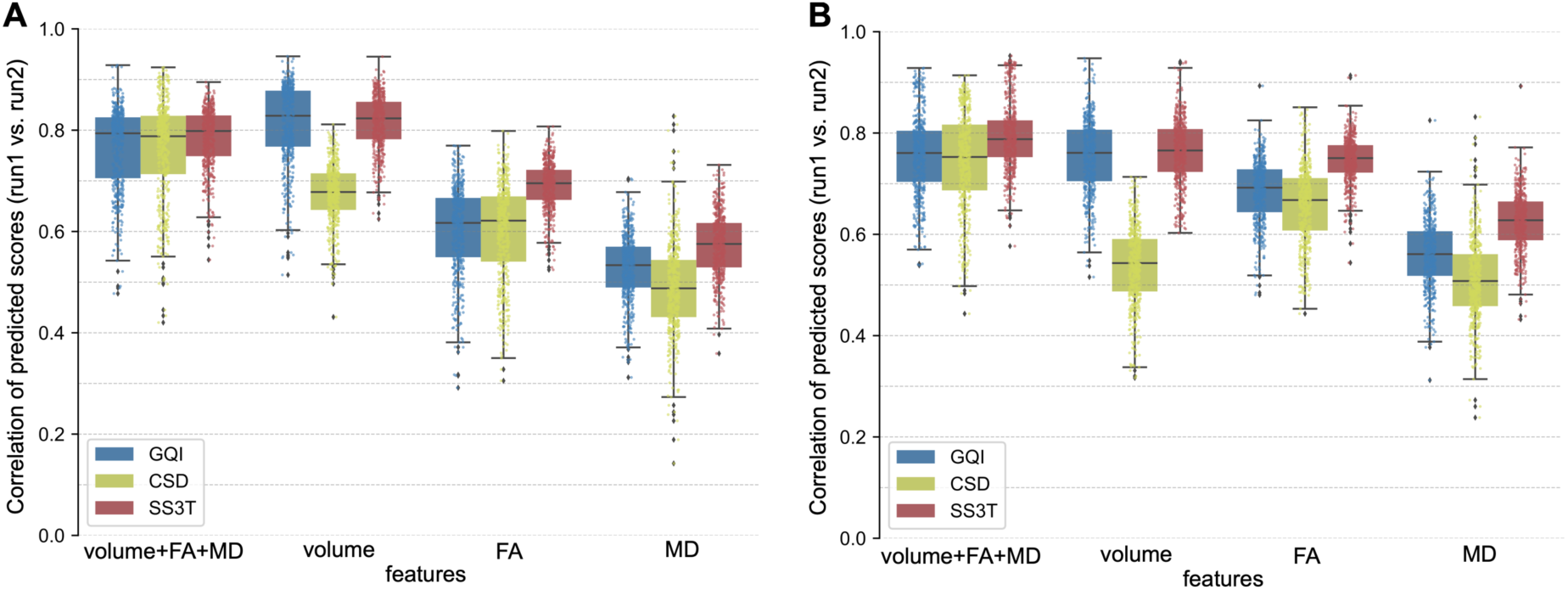
Comparison of the prediction reliability between different reconstruction methods for different groups of features for **A** predicting a composite IQ score and **B** predicting executive functioning. Prediction reliability was assessed by correlating the predictions obtained from features extracted from run-01 scans with predictions based on run-02 features for each fold. A higher correlation indicates a higher similarity and therefore also reliability across scans.

**Figure S7:**
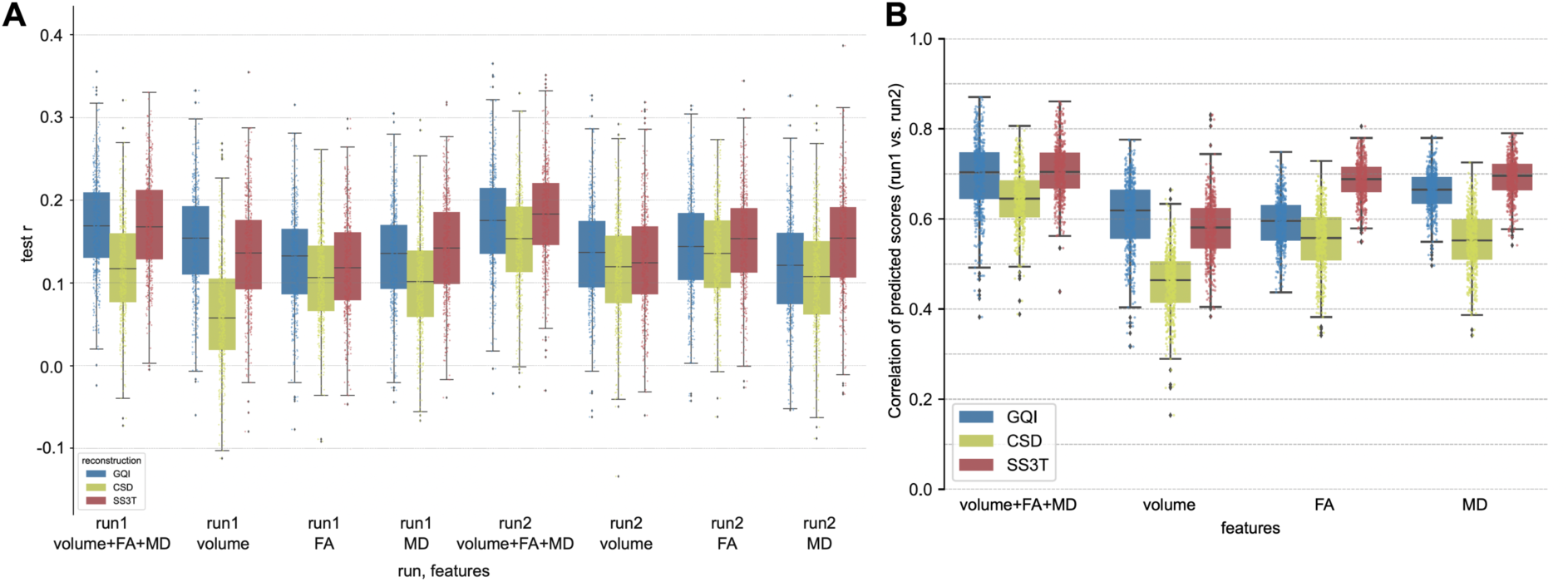
Prediction accuracy (**A**) and prediction reliability (**B**) when including TBV as a confound for predicting complex reasoning.

**Table S1:**
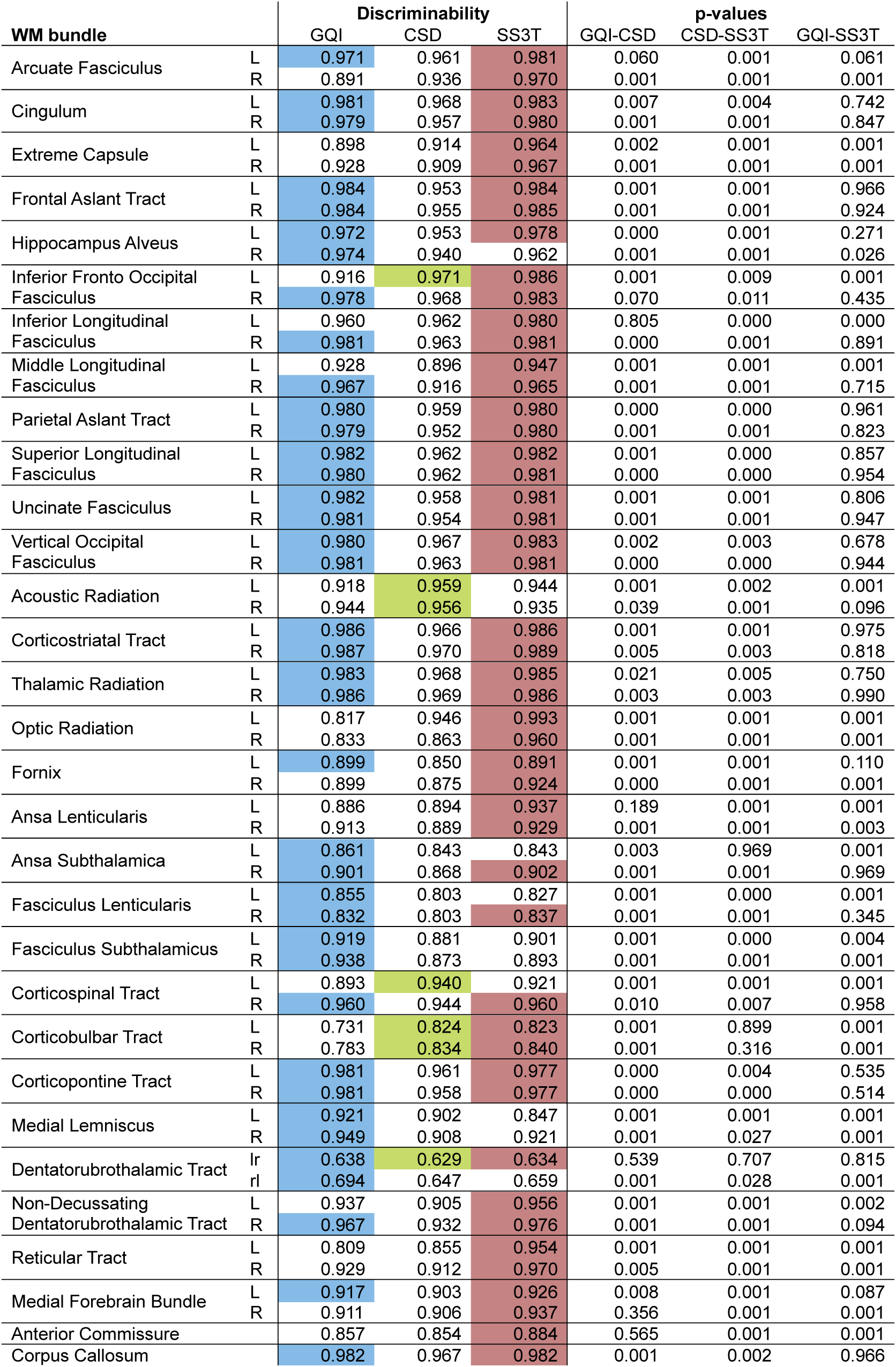
Discriminability per bundle per method including p-values. A single colored cell per row symbolizes that the corresponding method led to the best result and was significantly better than the other methods. Two or three colored cells per bundle show that there was no significant difference between the best two or three methods. Cells are colored according to the ODF reconstruction method (GQI: blue, CSD: green, SS3T: red) P-values are calculated using permutation tests as implemented in hyppo (Panda et al., 2020).

**Table S2:**
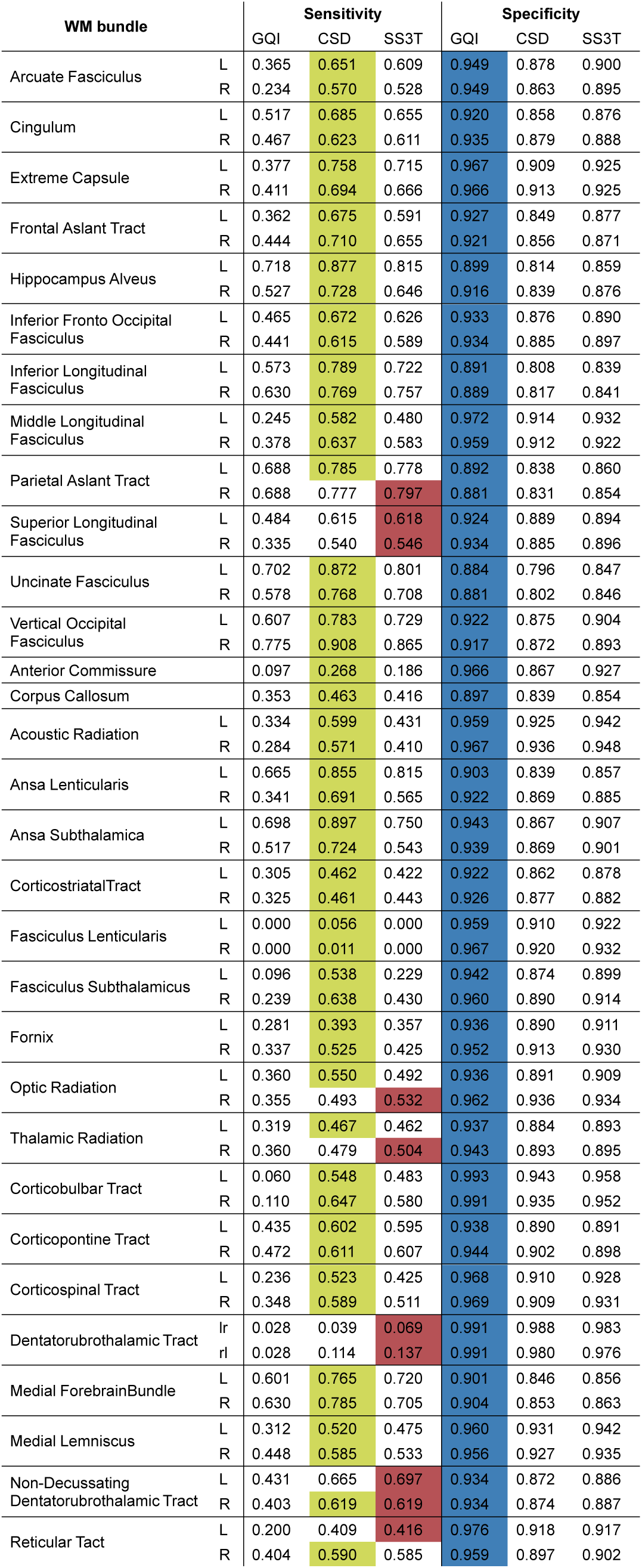
Median sensitivity and specificity per bundle and reconstruction method. The highest median sensitivity and specificity values are highlighted in red (GQI), green (CSD), or blue (SS3T), depending on the method with which they were achieved.

**Table S3:**
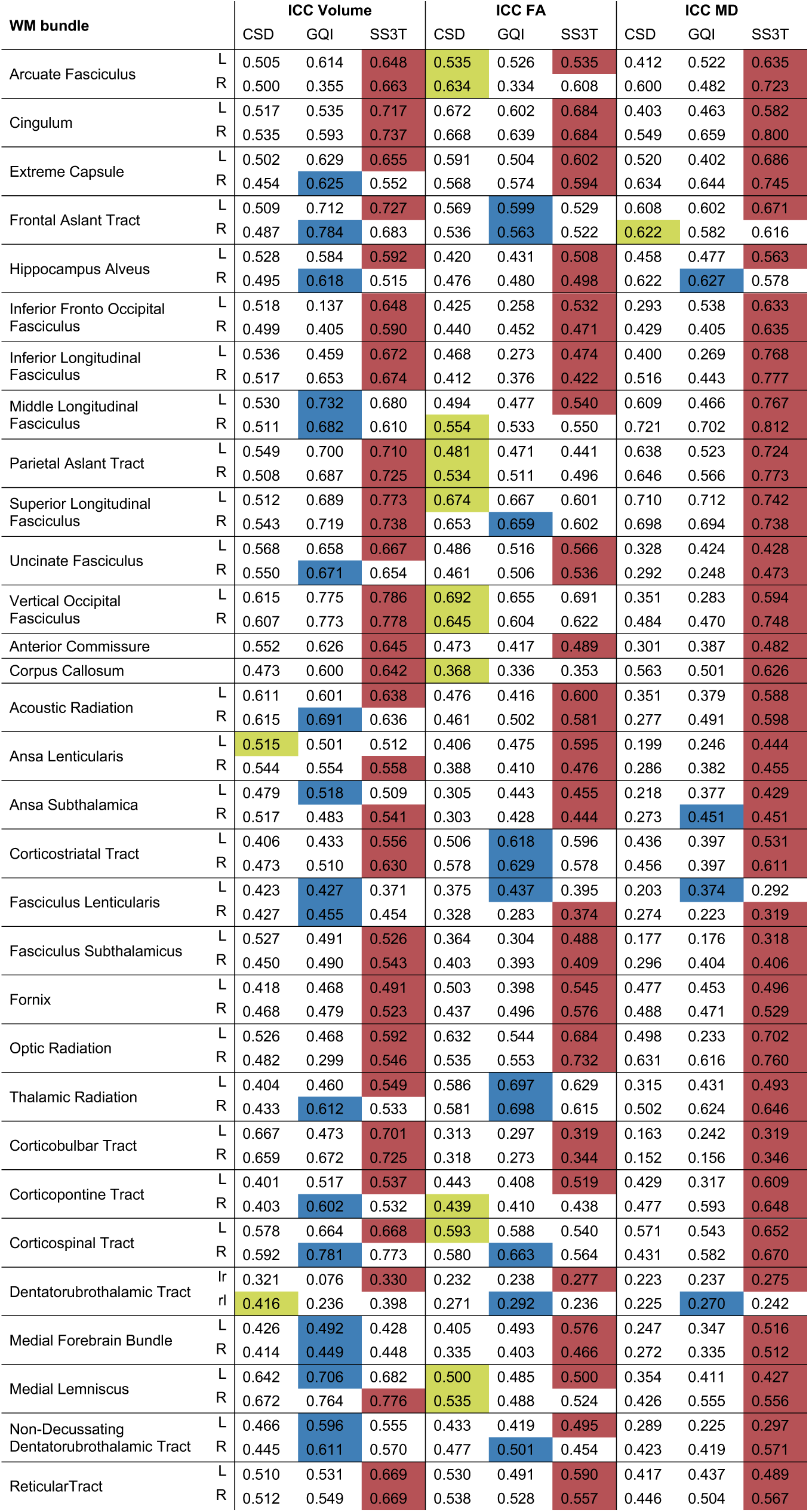
ICCs per bundle and reconstruction method for bundle volume, mean FA and mean MD. The highest ICCs for each bundle and each feature are highlighted in red (GQI), green (CSD), or blue (SS3T), depending on the method with which they were achieved.

