## Supplementary for "Benchmarking Orientation Distribution Function Estimation Methods for Tractometry in Single-Shell Diffusion Magnetic Resonance Imaging - An Evaluation of Test-Retest Reliability and Predictive Capability"

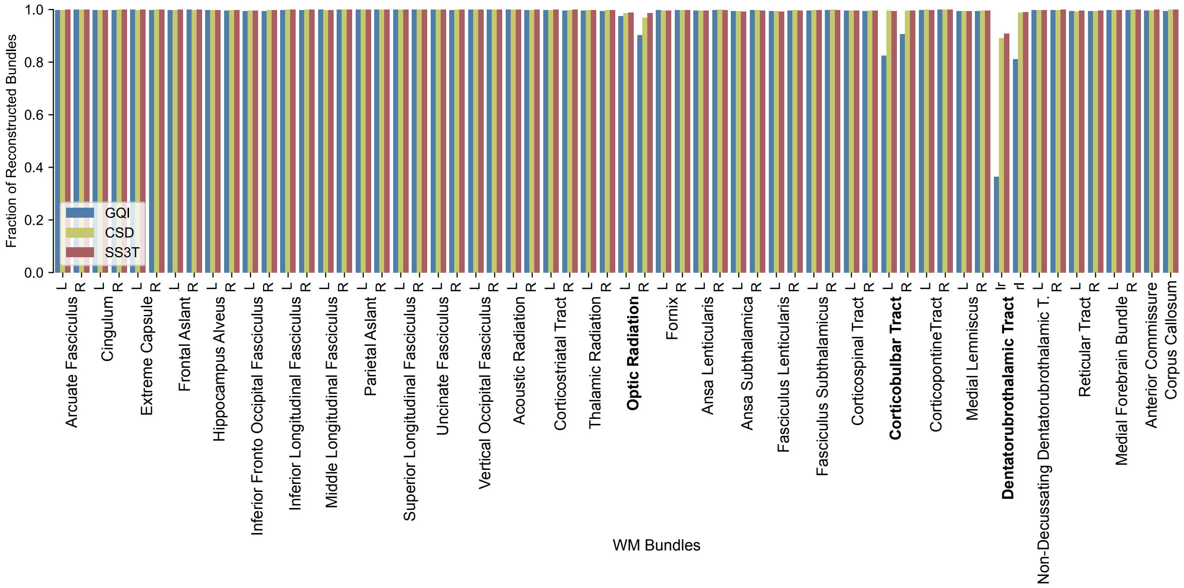


**Figure S1:** Reconstruction success rates per bundle per method. The reconstruction success rates represent the fraction of scans for which a given bundle could be reconstructed. Marked in bold are bundles with reconstruction fractions < 1 for at least one of the three reconstruction methods.

| WM bundle | | Discriminability GQI | CSD | SS3T | p-values GQI-CSD | CSD-SS3T | GQI-SS3T |
|---|---|---|---|---|---|---|---|
| Arcuate Fasciculus | L | 0.971 | 0.961 | 0.981 | 0.060 | 0.001 | 0.061 |
| | R | 0.891 | 0.936 | 0.970 | 0.001 | 0.001 | 0.001 |
| Cingulum | L | 0.981 | 0.968 | 0.983 | 0.007 | 0.004 | 0.742 |
| | R | 0.979 | 0.957 | 0.980 | 0.001 | 0.001 | 0.847 |
| Extreme Capsule | L | 0.898 | 0.914 | 0.964 | 0.002 | 0.001 | 0.001 |
| | R | 0.928 | 0.909 | 0.967 | 0.001 | 0.001 | 0.001 |
| Frontal Aslant Tract | L | 0.984 | 0.953 | 0.984 | 0.001 | 0.001 | 0.966 |
| | R | 0.984 | 0.955 | 0.985 | 0.001 | 0.001 | 0.924 |
| Hippocampus Alveus | L | 0.972 | 0.953 | 0.978 | 0.000 | 0.001 | 0.271 |
| | R | 0.974 | 0.940 | 0.962 | 0.001 | 0.001 | 0.026 |
| Inferior Fronto Occipital Fasciculus | L | 0.916 | 0.971 | 0.986 | 0.001 | 0.009 | 0.001 |
| | R | 0.978 | 0.968 | 0.983 | 0.070 | 0.011 | 0.435 |
| Inferior Longitudinal Fasciculus | L | 0.960 | 0.962 | 0.980 | 0.805 | 0.000 | 0.000 |
| | R | 0.981 | 0.963 | 0.981 | 0.000 | 0.001 | 0.891 |
| Middle Longitudinal Fasciculus | L | 0.928 | 0.896 | 0.947 | 0.001 | 0.001 | 0.001 |
| | R | 0.967 | 0.916 | 0.965 | 0.001 | 0.001 | 0.715 |
| Parietal Aslant Tract | L | 0.980 | 0.959 | 0.980 | 0.000 | 0.000 | 0.961 |
| | R | 0.979 | 0.952 | 0.980 | 0.001 | 0.001 | 0.823 |
| Superior Longitudinal Fasciculus | L | 0.982 | 0.962 | 0.982 | 0.001 | 0.000 | 0.857 |
| | R | 0.980 | 0.962 | 0.981 | 0.000 | 0.000 | 0.954 |
| Uncinate Fasciculus | L | 0.982 | 0.958 | 0.981 | 0.001 | 0.001 | 0.806 |
| | R | 0.981 | 0.954 | 0.981 | 0.001 | 0.001 | 0.947 |
| Vertical Occipital Fasciculus | L | 0.980 | 0.967 | 0.983 | 0.002 | 0.003 | 0.678 |
| | R | 0.981 | 0.963 | 0.981 | 0.000 | 0.000 | 0.944 |
| Acoustic Radiation | L | 0.918 | 0.959 | 0.944 | 0.001 | 0.002 | 0.001 |
| | R | 0.944 | 0.956 | 0.935 | 0.039 | 0.001 | 0.096 |
| Corticostriatal Tract | L | 0.986 | 0.966 | 0.986 | 0.001 | 0.001 | 0.975 |
| | R | 0.987 | 0.970 | 0.989 | 0.005 | 0.003 | 0.818 |
| Thalamic Radiation | L | 0.983 | 0.968 | 0.985 | 0.021 | 0.005 | 0.750 |
| | R | 0.986 | 0.969 | 0.986 | 0.003 | 0.003 | 0.990 |
| Optic Radiation | L | 0.817 | 0.946 | 0.993 | 0.001 | 0.001 | 0.001 |
| | R | 0.833 | 0.863 | 0.960 | 0.001 | 0.001 | 0.001 |
| Fornix | L | 0.899 | 0.850 | 0.891 | 0.001 | 0.001 | 0.110 |
| | R | 0.899 | 0.875 | 0.924 | 0.000 | 0.001 | 0.001 |
| Ansa Lenticularis | L | 0.886 | 0.894 | 0.937 | 0.189 | 0.001 | 0.001 |
| | R | 0.913 | 0.889 | 0.929 | 0.001 | 0.001 | 0.003 |
| Ansa Subthalamica | L | 0.861 | 0.843 | 0.843 | 0.003 | 0.969 | 0.001 |
| | R | 0.901 | 0.868 | 0.902 | 0.001 | 0.001 | 0.969 |
| Fasciculus Lenticularis | L | 0.855 | 0.803 | 0.827 | 0.001 | 0.000 | 0.001 |
| | R | 0.832 | 0.803 | 0.837 | 0.001 | 0.001 | 0.345 |
| Fasciculus Subthalamicus | L | 0.919 | 0.881 | 0.901 | 0.001 | 0.000 | 0.004 |
| | R | 0.938 | 0.873 | 0.893 | 0.001 | 0.001 | 0.001 |
| Corticospinal Tract | L | 0.893 | 0.940 | 0.921 | 0.001 | 0.001 | 0.001 |
| | R | 0.960 | 0.944 | 0.960 | 0.010 | 0.007 | 0.958 |
| Corticobulbar Tract | L | 0.731 | 0.824 | 0.823 | 0.001 | 0.899 | 0.001 |
| | R | 0.783 | 0.834 | 0.840 | 0.001 | 0.316 | 0.001 |
| Corticopontine Tract | L | 0.981 | 0.961 | 0.977 | 0.000 | 0.004 | 0.535 |
| | R | 0.981 | 0.958 | 0.977 | 0.000 | 0.000 | 0.514 |
| Medial Lemniscus | L | 0.921 | 0.902 | 0.847 | 0.001 | 0.001 | 0.001 |
| | R | 0.949 | 0.908 | 0.921 | 0.001 | 0.027 | 0.001 |
| Dentatorubrothalamic Tract | lr | 0.638 | 0.629 | 0.634 | 0.539 | 0.707 | 0.815 |
| | rl | 0.694 | 0.647 | 0.659 | 0.001 | 0.028 | 0.001 |
| Non-Decussating Dentatorubrothalamic Tract | L | 0.937 | 0.905 | 0.956 | 0.001 | 0.001 | 0.002 |
| | R | 0.967 | 0.932 | 0.976 | 0.001 | 0.001 | 0.094 |
| Reticular Tract | L | 0.809 | 0.855 | 0.954 | 0.001 | 0.001 | 0.001 |
| | R | 0.929 | 0.912 | 0.970 | 0.005 | 0.001 | 0.001 |
| Medial Forebrain Bundle | L | 0.917 | 0.903 | 0.926 | 0.008 | 0.001 | 0.087 |
| | R | 0.911 | 0.906 | 0.937 | 0.356 | 0.001 | 0.001 |
| Anterior Commissure | | 0.857 | 0.854 | 0.884 | 0.565 | 0.001 | 0.001 |
| Corpus Callosum | | 0.982 | 0.967 | 0.982 | 0.001 | 0.002 | 0.966 |

**Table S1:** Discriminability per bundle per method including p-values. A single colored cell per row symbolizes that the corresponding method led to the best result and was significantly better than the other methods. Two or three colored cells per bundle show that there was no significant difference between the best two or three methods. Cells are colored according to the ODF reconstruction method (GQI: blue, CSD: green, SS3T: red) P-values are calculated using permutation tests as implemented in hyppo (Panda et al., 2020).

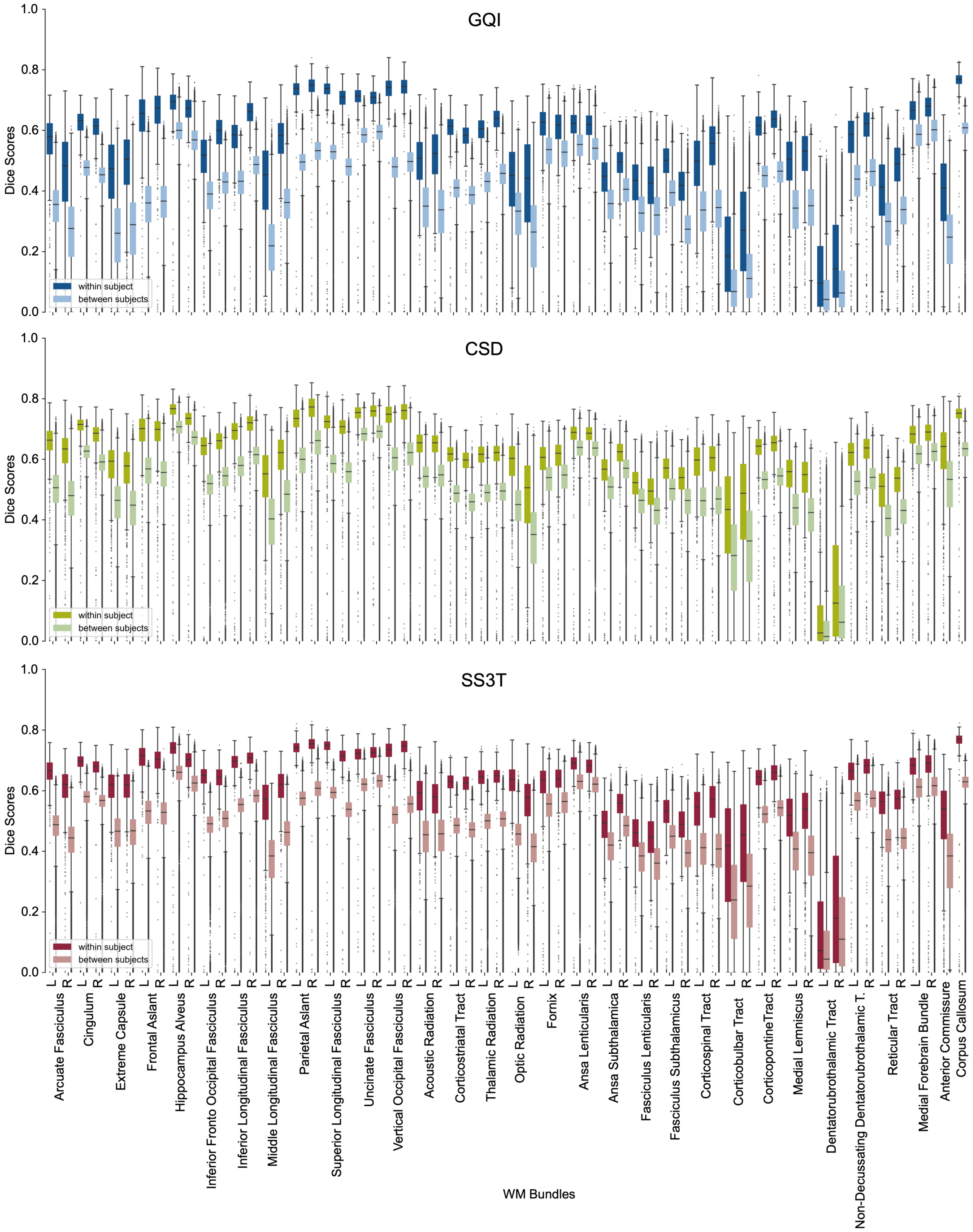


**Figure S2:** Full distributions of within and between-subject dice scores per bundle for GQI, CSD, and SS3T.

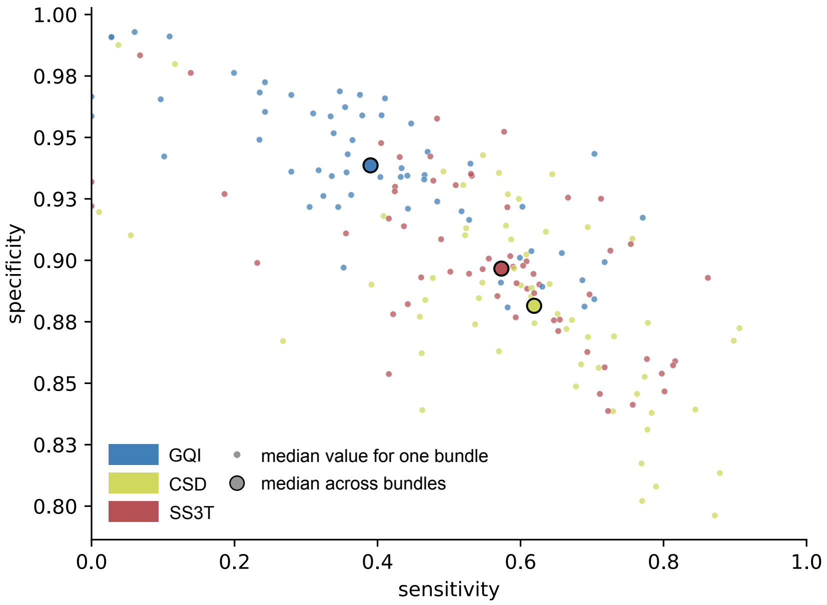


**Figure S3:** Median sensitivity and specificity for all 60 reconstructed WM bundles. Sensitivity and specificity were calculated for each instance (different subjects, different scans) of the reconstructed bundle. Visualized here are the median sensitivity and specificity values across all instances of a given bundle.

| WM bundle | | Sensitivity | | | Specificity | | |
|---|---|---|---|---|---|---|---|
| | | GQI | CSD | SS3T | GQI | CSD | SS3T |
| Arcuate Fasciculus | L | 0.365 | 0.651 | 0.609 | 0.949 | 0.878 | 0.900 |
| | R | 0.234 | 0.570 | 0.528 | 0.949 | 0.863 | 0.895 |
| Cingulum | L | 0.517 | 0.685 | 0.655 | 0.920 | 0.858 | 0.876 |
| | R | 0.467 | 0.623 | 0.611 | 0.935 | 0.879 | 0.888 |
| Extreme Capsule | L | 0.377 | 0.758 | 0.715 | 0.967 | 0.909 | 0.925 |
| | R | 0.411 | 0.694 | 0.666 | 0.966 | 0.913 | 0.925 |
| Frontal Aslant Tract | L | 0.362 | 0.675 | 0.591 | 0.927 | 0.849 | 0.877 |
| | R | 0.444 | 0.710 | 0.655 | 0.921 | 0.856 | 0.871 |
| Hippocampus Alveus | L | 0.718 | 0.877 | 0.815 | 0.899 | 0.814 | 0.859 |
| | R | 0.527 | 0.728 | 0.646 | 0.916 | 0.839 | 0.876 |
| Inferior Fronto Occipital Fasciculus | L | 0.465 | 0.672 | 0.626 | 0.933 | 0.876 | 0.890 |
| | R | 0.441 | 0.615 | 0.589 | 0.934 | 0.885 | 0.897 |
| Inferior Longitudinal Fasciculus | L | 0.573 | 0.789 | 0.722 | 0.891 | 0.808 | 0.839 |
| | R | 0.630 | 0.769 | 0.757 | 0.889 | 0.817 | 0.841 |
| Middle Longitudinal Fasciculus | L | 0.245 | 0.582 | 0.480 | 0.972 | 0.914 | 0.932 |
| | R | 0.378 | 0.637 | 0.583 | 0.959 | 0.912 | 0.922 |
| Parietal Aslant Tract | L | 0.688 | 0.785 | 0.778 | 0.892 | 0.838 | 0.860 |
| | R | 0.688 | 0.777 | 0.797 | 0.881 | 0.831 | 0.854 |
| Superior Longitudinal Fasciculus | L | 0.484 | 0.615 | 0.618 | 0.924 | 0.889 | 0.894 |
| | R | 0.335 | 0.540 | 0.546 | 0.934 | 0.885 | 0.896 |
| Uncinate Fasciculus | L | 0.702 | 0.872 | 0.801 | 0.884 | 0.796 | 0.847 |
| | R | 0.578 | 0.768 | 0.708 | 0.881 | 0.802 | 0.846 |
| Vertical Occipital Fasciculus | L | 0.607 | 0.783 | 0.729 | 0.922 | 0.875 | 0.904 |
| | R | 0.775 | 0.908 | 0.865 | 0.917 | 0.872 | 0.893 |
| Anterior Commissure | | 0.097 | 0.268 | 0.186 | 0.966 | 0.867 | 0.927 |
| Corpus Callosum | | 0.353 | 0.463 | 0.416 | 0.897 | 0.839 | 0.854 |
| Acoustic Radiation | L | 0.334 | 0.599 | 0.431 | 0.959 | 0.925 | 0.942 |
| | R | 0.284 | 0.571 | 0.410 | 0.967 | 0.936 | 0.948 |
| Ansa Lenticularis | L | 0.665 | 0.855 | 0.815 | 0.903 | 0.839 | 0.857 |
| | R | 0.341 | 0.691 | 0.565 | 0.922 | 0.869 | 0.885 |
| Ansa Subthalamica | L | 0.698 | 0.897 | 0.750 | 0.943 | 0.867 | 0.907 |
| | R | 0.517 | 0.724 | 0.543 | 0.939 | 0.869 | 0.901 |
| CorticostriatalTract | L | 0.305 | 0.462 | 0.422 | 0.922 | 0.862 | 0.878 |
| | R | 0.325 | 0.461 | 0.443 | 0.926 | 0.877 | 0.882 |
| Fasciculus Lenticularis | L | 0.000 | 0.056 | 0.000 | 0.959 | 0.910 | 0.922 |
| | R | 0.000 | 0.011 | 0.000 | 0.967 | 0.920 | 0.932 |
| Fasciculus Subthalamicus | L | 0.096 | 0.538 | 0.229 | 0.942 | 0.874 | 0.899 |
| | R | 0.239 | 0.638 | 0.430 | 0.960 | 0.890 | 0.914 |
| Fornix | L | 0.281 | 0.393 | 0.357 | 0.936 | 0.890 | 0.911 |
| | R | 0.337 | 0.525 | 0.425 | 0.952 | 0.913 | 0.930 |
| Optic Radiation | L | 0.360 | 0.550 | 0.492 | 0.936 | 0.891 | 0.909 |
| | R | 0.355 | 0.493 | 0.532 | 0.962 | 0.936 | 0.934 |
| Thalamic Radiation | L | 0.319 | 0.467 | 0.462 | 0.937 | 0.884 | 0.893 |
| | R | 0.360 | 0.479 | 0.504 | 0.943 | 0.893 | 0.895 |
| Corticobulbar Tract | L | 0.060 | 0.548 | 0.483 | 0.993 | 0.943 | 0.958 |
| | R | 0.110 | 0.647 | 0.580 | 0.991 | 0.935 | 0.952 |
| Corticopontine Tract | L | 0.435 | 0.602 | 0.595 | 0.938 | 0.890 | 0.891 |
| | R | 0.472 | 0.611 | 0.607 | 0.944 | 0.902 | 0.898 |
| Corticospinal Tract | L | 0.236 | 0.523 | 0.425 | 0.968 | 0.910 | 0.928 |
| | R | 0.348 | 0.589 | 0.511 | 0.969 | 0.909 | 0.931 |
| Dentatorubrothalamic Tract | lr | 0.028 | 0.039 | 0.069 | 0.991 | 0.988 | 0.983 |
| | rl | 0.028 | 0.114 | 0.137 | 0.991 | 0.980 | 0.976 |
| Medial ForebrainBundle | L | 0.601 | 0.765 | 0.720 | 0.901 | 0.846 | 0.856 |
| | R | 0.630 | 0.785 | 0.705 | 0.904 | 0.853 | 0.863 |
| Medial Lemniscus | L | 0.312 | 0.520 | 0.475 | 0.960 | 0.931 | 0.942 |
| | R | 0.448 | 0.585 | 0.533 | 0.956 | 0.927 | 0.935 |
| Non-Decussating Dentatorubrothalamic Tract | L | 0.431 | 0.665 | 0.697 | 0.934 | 0.872 | 0.886 |
| | R | 0.403 | 0.619 | 0.619 | 0.934 | 0.874 | 0.887 |
| Reticular Tact | L | 0.200 | 0.409 | 0.416 | 0.976 | 0.918 | 0.917 |
| | R | 0.404 | 0.590 | 0.585 | 0.959 | 0.897 | 0.902 |

**Table S2:** Median sensitivity and specificity per bundle and reconstruction method. The highest median sensitivity and specificity values are highlighted in red (GQI), green (CSD), or blue (SS3T), depending on the method with which they were achieved.

| WM bundle | | ICC Volume | | | ICC FA | | | ICC MD | | |
|---|---|---|---|---|---|---|---|---|---|---|
| | | CSD | GQI | SS3T | CSD | GQI | SS3T | CSD | GQI | SS3T |
| Arcuate Fasciculus | L | 0.505 | 0.614 | 0.648 | 0.535 | 0.526 | 0.535 | 0.412 | 0.522 | 0.635 |
| | R | 0.500 | 0.355 | 0.663 | 0.634 | 0.334 | 0.608 | 0.600 | 0.482 | 0.723 |
| Cingulum | L | 0.517 | 0.535 | 0.717 | 0.672 | 0.602 | 0.684 | 0.403 | 0.463 | 0.582 |
| | R | 0.535 | 0.593 | 0.737 | 0.668 | 0.639 | 0.684 | 0.549 | 0.659 | 0.800 |
| Extreme Capsule | L | 0.502 | 0.629 | 0.655 | 0.591 | 0.504 | 0.602 | 0.520 | 0.402 | 0.686 |
| | R | 0.454 | 0.625 | 0.552 | 0.568 | 0.574 | 0.594 | 0.634 | 0.644 | 0.745 |
| Frontal Aslant Tract | L | 0.509 | 0.712 | 0.727 | 0.569 | 0.599 | 0.529 | 0.608 | 0.602 | 0.671 |
| | R | 0.487 | 0.784 | 0.683 | 0.536 | 0.563 | 0.522 | 0.622 | 0.582 | 0.616 |
| Hippocampus Alveus | L | 0.528 | 0.584 | 0.592 | 0.420 | 0.431 | 0.508 | 0.458 | 0.477 | 0.563 |
| | R | 0.495 | 0.618 | 0.515 | 0.476 | 0.480 | 0.498 | 0.622 | 0.627 | 0.578 |
| Inferior Fronto Occipital Fasciculus | L | 0.518 | 0.137 | 0.648 | 0.425 | 0.258 | 0.532 | 0.293 | 0.538 | 0.633 |
| | R | 0.499 | 0.405 | 0.590 | 0.440 | 0.452 | 0.471 | 0.429 | 0.405 | 0.635 |
| Inferior Longitudinal Fasciculus | L | 0.536 | 0.459 | 0.672 | 0.468 | 0.273 | 0.474 | 0.400 | 0.269 | 0.768 |
| | R | 0.517 | 0.653 | 0.674 | 0.412 | 0.376 | 0.422 | 0.516 | 0.443 | 0.777 |
| Middle Longitudinal Fasciculus | L | 0.530 | 0.732 | 0.680 | 0.494 | 0.477 | 0.540 | 0.609 | 0.466 | 0.767 |
| | R | 0.511 | 0.682 | 0.610 | 0.554 | 0.533 | 0.550 | 0.721 | 0.702 | 0.812 |
| Parietal Aslant Tract | L | 0.549 | 0.700 | 0.710 | 0.481 | 0.471 | 0.441 | 0.638 | 0.523 | 0.724 |
| | R | 0.508 | 0.687 | 0.725 | 0.534 | 0.511 | 0.496 | 0.646 | 0.566 | 0.773 |
| Superior Longitudinal Fasciculus | L | 0.512 | 0.689 | 0.773 | 0.674 | 0.667 | 0.601 | 0.710 | 0.712 | 0.742 |
| | R | 0.543 | 0.719 | 0.738 | 0.653 | 0.659 | 0.602 | 0.698 | 0.694 | 0.738 |
| Uncinate Fasciculus | L | 0.568 | 0.658 | 0.667 | 0.486 | 0.516 | 0.566 | 0.328 | 0.424 | 0.428 |
| | R | 0.550 | 0.671 | 0.654 | 0.461 | 0.506 | 0.536 | 0.292 | 0.248 | 0.473 |
| Vertical Occipital Fasciculus | L | 0.615 | 0.775 | 0.786 | 0.692 | 0.655 | 0.691 | 0.351 | 0.283 | 0.594 |
| | R | 0.607 | 0.773 | 0.778 | 0.645 | 0.604 | 0.622 | 0.484 | 0.470 | 0.748 |
| Anterior Commissure | | 0.552 | 0.626 | 0.645 | 0.473 | 0.417 | 0.489 | 0.301 | 0.387 | 0.482 |
| Corpus Callosum | | 0.473 | 0.600 | 0.642 | 0.368 | 0.336 | 0.353 | 0.563 | 0.501 | 0.626 |
| Acoustic Radiation | L | 0.611 | 0.601 | 0.638 | 0.476 | 0.416 | 0.600 | 0.351 | 0.379 | 0.588 |
| | R | 0.615 | 0.691 | 0.636 | 0.461 | 0.502 | 0.581 | 0.277 | 0.491 | 0.598 |
| Ansa Lenticularis | L | 0.515 | 0.501 | 0.512 | 0.406 | 0.475 | 0.595 | 0.199 | 0.246 | 0.444 |
| | R | 0.544 | 0.554 | 0.558 | 0.388 | 0.410 | 0.476 | 0.286 | 0.382 | 0.455 |
| Ansa Subthalamica | L | 0.479 | 0.518 | 0.509 | 0.305 | 0.443 | 0.455 | 0.218 | 0.377 | 0.429 |
| | R | 0.517 | 0.483 | 0.541 | 0.303 | 0.428 | 0.444 | 0.273 | 0.451 | 0.451 |
| Corticostriatal Tract | L | 0.406 | 0.433 | 0.556 | 0.506 | 0.618 | 0.596 | 0.436 | 0.397 | 0.531 |
| | R | 0.473 | 0.510 | 0.630 | 0.578 | 0.629 | 0.578 | 0.456 | 0.397 | 0.611 |
| Fasciculus Lenticularis | L | 0.423 | 0.427 | 0.371 | 0.375 | 0.437 | 0.395 | 0.203 | 0.374 | 0.292 |
| | R | 0.427 | 0.455 | 0.454 | 0.328 | 0.283 | 0.374 | 0.274 | 0.223 | 0.319 |
| Fasciculus Subthalamicus | L | 0.527 | 0.491 | 0.526 | 0.364 | 0.304 | 0.488 | 0.177 | 0.176 | 0.318 |
| | R | 0.450 | 0.490 | 0.543 | 0.403 | 0.393 | 0.409 | 0.296 | 0.404 | 0.406 |
| Fornix | L | 0.418 | 0.468 | 0.491 | 0.503 | 0.398 | 0.545 | 0.477 | 0.453 | 0.496 |
| | R | 0.468 | 0.479 | 0.523 | 0.437 | 0.496 | 0.576 | 0.488 | 0.471 | 0.529 |
| Optic Radiation | L | 0.526 | 0.468 | 0.592 | 0.632 | 0.544 | 0.684 | 0.498 | 0.233 | 0.702 |
| | R | 0.482 | 0.299 | 0.546 | 0.535 | 0.553 | 0.732 | 0.631 | 0.616 | 0.760 |
| Thalamic Radiation | L | 0.404 | 0.460 | 0.549 | 0.586 | 0.697 | 0.629 | 0.315 | 0.431 | 0.493 |
| | R | 0.433 | 0.612 | 0.533 | 0.581 | 0.698 | 0.615 | 0.502 | 0.624 | 0.646 |
| Corticobulbar Tract | L | 0.667 | 0.473 | 0.701 | 0.313 | 0.297 | 0.319 | 0.163 | 0.242 | 0.319 |
| | R | 0.659 | 0.672 | 0.725 | 0.318 | 0.273 | 0.344 | 0.152 | 0.156 | 0.346 |
| Corticopontine Tract | L | 0.401 | 0.517 | 0.537 | 0.443 | 0.408 | 0.519 | 0.429 | 0.317 | 0.609 |
| | R | 0.403 | 0.602 | 0.532 | 0.439 | 0.410 | 0.438 | 0.477 | 0.593 | 0.648 |
| Corticospinal Tract | L | 0.578 | 0.664 | 0.668 | 0.593 | 0.588 | 0.540 | 0.571 | 0.543 | 0.652 |
| | R | 0.592 | 0.781 | 0.773 | 0.580 | 0.663 | 0.564 | 0.431 | 0.582 | 0.670 |
| Dentatorubrothalamic Tract | lr | 0.321 | 0.076 | 0.330 | 0.232 | 0.238 | 0.277 | 0.223 | 0.237 | 0.275 |
| | rl | 0.416 | 0.236 | 0.398 | 0.271 | 0.292 | 0.236 | 0.225 | 0.270 | 0.242 |
| Medial Forebrain Bundle | L | 0.426 | 0.492 | 0.428 | 0.405 | 0.493 | 0.576 | 0.247 | 0.347 | 0.516 |
| | R | 0.414 | 0.449 | 0.448 | 0.335 | 0.403 | 0.466 | 0.272 | 0.335 | 0.512 |
| Medial Lemniscus | L | 0.642 | 0.706 | 0.682 | 0.500 | 0.485 | 0.500 | 0.354 | 0.411 | 0.427 |
| | R | 0.672 | 0.764 | 0.776 | 0.535 | 0.488 | 0.524 | 0.426 | 0.555 | 0.556 |
| Non-Decussating Dentatorubrothalamic Tract | L | 0.466 | 0.596 | 0.555 | 0.433 | 0.419 | 0.495 | 0.289 | 0.225 | 0.297 |
| | R | 0.445 | 0.611 | 0.570 | 0.477 | 0.501 | 0.454 | 0.423 | 0.419 | 0.571 |
| ReticularTract | L | 0.510 | 0.531 | 0.669 | 0.530 | 0.491 | 0.590 | 0.417 | 0.437 | 0.489 |
| | R | 0.512 | 0.549 | 0.669 | 0.538 | 0.528 | 0.557 | 0.446 | 0.504 | 0.567 |

**Table S3:** ICCs per bundle and reconstruction method for bundle volume, mean FA and mean MD. The highest ICCs for each bundle and each feature are highlighted in red (GQI), green (CSD), or blue (SS3T), depending on the method with which they were achieved.

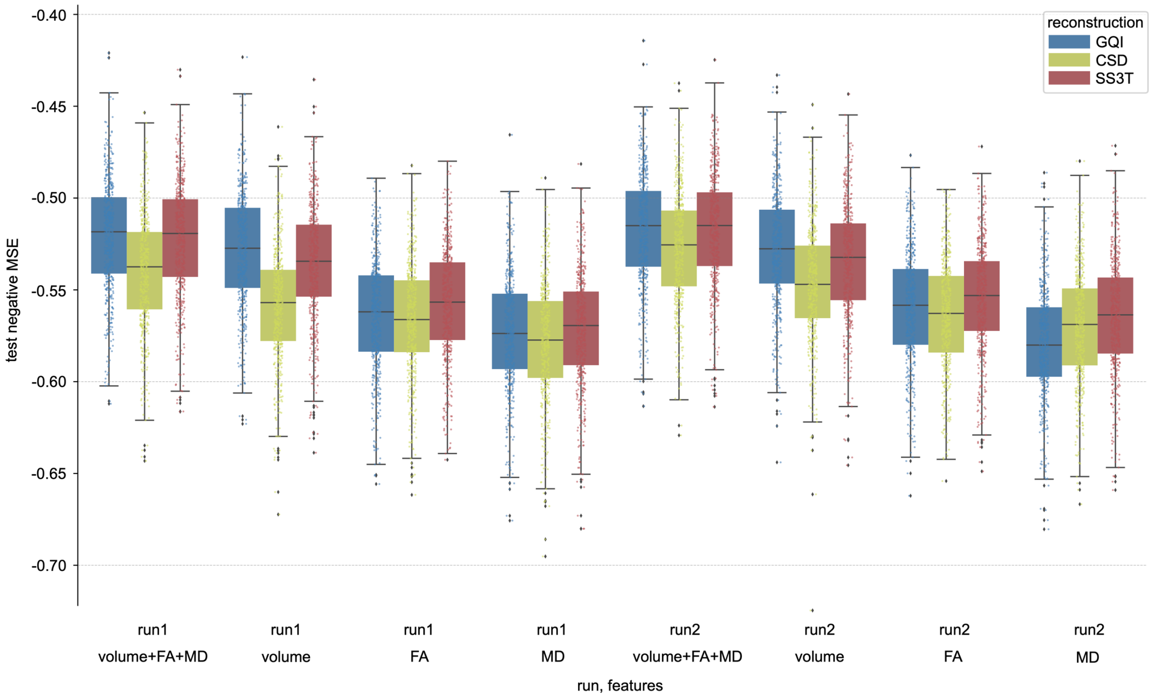


**Figure S4:** Prediction accuracy in terms of negative MSE for predicting complex reasoning from different groups of features for all three reconstruction methods. Each distribution contains 500 points (100 x 5-fold CV). The left block of prediction accuracies used features extracted from bundles reconstructed from run-01 scans, the right block from run-02 scans. For each run, four different groups of features were evaluated: volume, mean FA and mean MD for each of the 54 considered bundles (162 features), only the bundle volume (54 features), only the mean FA (54 features), and only the mean MD (54 features).

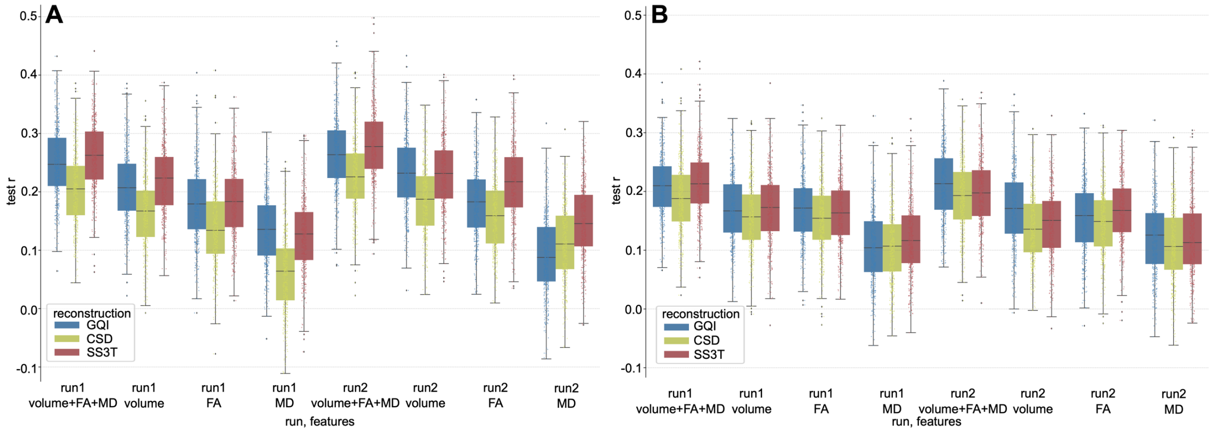


**Figure S5:** Prediction accuracy in terms of Pearson correlation for predicting **A** a composite IQ score and **B** executive functioning from different groups of features for all three reconstruction methods. Each distribution contains 500 points (100 x 5-fold CV). The left block of prediction accuracies used features extracted from bundles reconstructed from run-01 scans, and the right block from run-02 scans. For each run, four different groups of features were evaluated: volume, mean FA, and mean MD for each of the 54 considered bundles (162 features), only the bundle volume (54 features), only the mean FA (54 features), and only the mean MD (54 features).

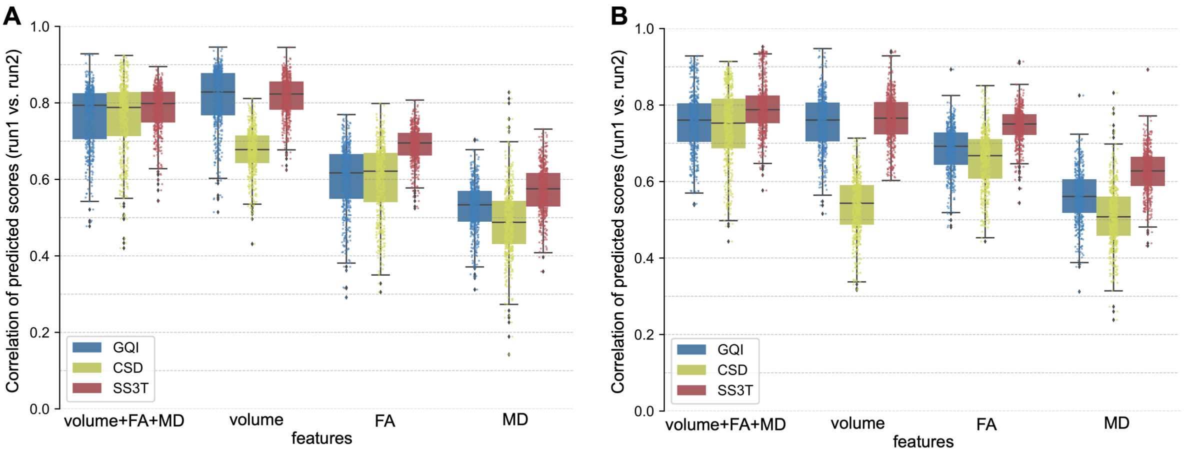


**Figure S6:** Comparison of the prediction reliability between different reconstruction methods for different groups of features for **A** predicting a composite IQ score and **B** predicting executive functioning. Prediction reliability was assessed by correlating the predictions obtained from features extracted from run-01 scans with predictions based on run-02 features for each fold. A higher correlation indicates a higher similarity and therefore also reliability across scans.

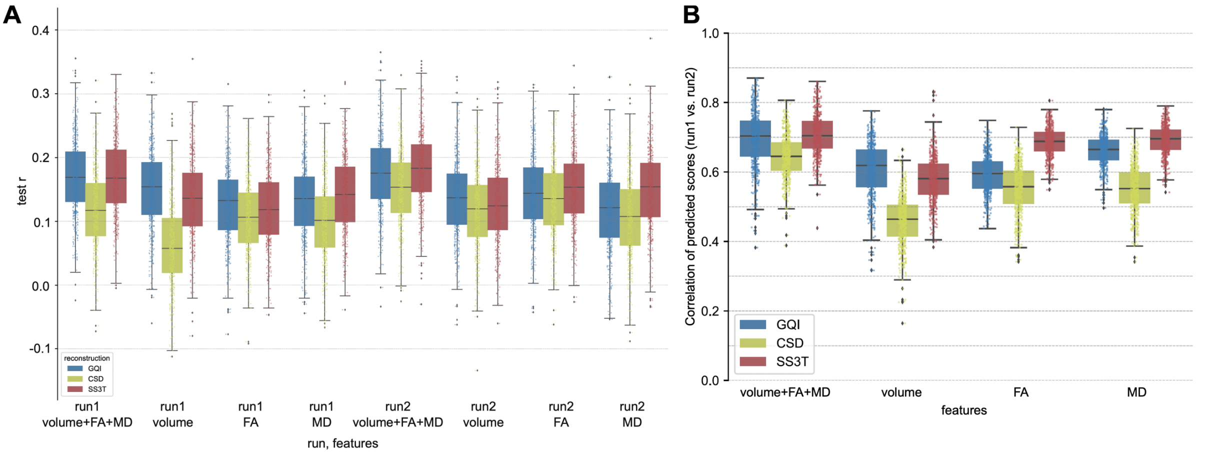


**Figure S7:** Prediction accuracy (**A**) and prediction reliability (**B**) when including TBV as a confound for predicting complex reasoning.
